# The m^6^A reader Ythdf restricts axonal growth in *Drosophila* through target selection modulation of the Fragile X mental retardation protein

**DOI:** 10.1101/2020.03.04.976886

**Authors:** Alessia Soldano, Lina Worpenberg, Chiara Paolantoni, Sara Longhi, Miriam M. Mulorz, Tina Lence, Hans-Hermann Wessels, Giuseppe Aiello, Michela Notarangelo, FX Reymond Sutandy, Marion Scheibe, Raghu R. Edupuganti, Anke Busch, Martin M. Möckel, Michiel Vermeulen, Falk Butter, Julian König, Uwe Ohler, Christoph Dieterich, Alessandro Quattrone, Jean-Yves Roignant

## Abstract

N6-methyladenosine (m^6^A) regulates a variety of physiological processes through modulation of RNA metabolism. The modification is particularly enriched in the nervous system of several species, and its dysregulation has been associated with neurodevelopmental defects and neural dysfunctions. In *Drosophila*, loss of m^6^A alters fly behavior albeit the underlying mechanism and the role of m^6^A during nervous system development have remained elusive. Here we find that impairment of the m^6^A pathway leads to axonal overgrowth and misguidance at larval neuromuscular junctions as well as in the adult mushroom bodies. We identify Ythdf as the main m^6^A reader in the nervous system being required for limiting axonal growth. Mechanistically, we show that Ythdf directly interacts with Fragile X mental retardation protein to inhibit the translation of key transcripts involved in axonal growth regulation. Altogether, this study demonstrates that the m^6^A pathway controls development of the nervous system by modulating Fmr1 target selection.

## Introduction

Chemical modifications on DNA and histones impact gene expression during cell differentiation, organismal development and in several other biological programs (Jaenisch and Bird, 2003). Similarly, RNA modifications represent a novel layer of gene regulation but their functional characterization during development and in other biological/pathological processes is still incomplete in its infancy (Hsu et al., 2017).

N6-methyladenosine (m^6^A) is the most prevalent modification found in mRNA and long non-coding RNA. The mark is widely conserved, enriched in mRNAs at the beginning of the last exons, in the sequence context RRACH (where R=A or G and H= A, C or U) (Dominissini et al., 2012; Garcia-Campos et al., 2019; Meyer et al., 2012). m^6^A plays a central role in modulating RNA function, since it can influence many aspects of RNA life such as splicing, export, translation and decay (Roignant and Soller, 2017; Zhao et al., 2017). m^6^A deposition is operated by a multi-subunit methyltransferase complex, composed of METTL3 and METTL14, as well as several associated components (reviewed in (Lence et al., 2019)) and can be reverted to adenosine by demethylases, such as FTO and ALKBH5 (Jia et al., 2011; Zheng et al., 2013). The m^6^A signature is subsequently recognized by “reader” proteins, among which the best described is the YTH domain family of proteins that decode the signal and mediate m^6^A biological effects (Liao et al., 2018; Patil et al., 2018).

Increasing evidence suggests a central role of m^6^A during nervous system development and functions (Angelova et al., 2018; Du et al., 2019; Jung and Goldman, 2018; Li et al., 2019; Livneh et al., 2020; Widagdo and Anggono, 2018). m^6^A is present at particularly high levels in the nervous system of different model animals (Lence et al., 2016; Meyer et al., 2012), and these levels can vary following behavioral stimuli or sensory experience (Engel et al., 2018; Koranda et al., 2018; Widagdo et al., 2016; Yoon et al., 2018). In mouse, m^6^A controls brain development (Chen et al., 2019; Li et al., 2017; Li et al., 2018; Ma et al., 2018; Wang et al., 2018; Zhuang et al., 2019) and is also required for axon regeneration (Weng et al., 2018) and synaptic functions (Engel et al., 2018; Koranda et al., 2018; Merkurjev et al., 2018; Shi et al., 2018; Yu et al., 2018). Similarly, in *Drosophila*, m^6^A promotes flight and locomotion via a neuronal function (Haussmann et al., 2016; Kan et al., 2017; Lence et al., 2016; Lence et al., 2017). A proper level of m^6^A appears critical for regulating axonal growth as *Mettl3* knock out (KO) in *Drosophila* has been associated with axonal overgrowth at neuromuscular junctions (NMJs) (Lence et al., 2016), and conversely, higher m^6^A (or m^6^Am) produced by the loss of FTO, leads to shorter axonal length in mouse dorsal root ganglia neurons (Yu et al., 2018). To date, the underlying mechanism of m^6^A in axonal growth has remained elusive.

Fragile X mental retardation protein (FMRP) is a polyribosome-associated RNA binding protein (RBP) that negatively regulates the translation of a subset of dendritic mRNAs (Darnell et al., 2011; Jacquemont et al., 2018; Laggerbauer et al., 2001; Li et al., 2001). Additional functions in splicing, export and mRNA stability have also been reported for this RBP (Davis and Broadie, 2017). The loss of FMRP is the genetic cause of Fragile X syndrome (FXS), the most common inherited form of intellectual disability and autism, with an estimated prevalence of 1 in 4000 males, and 1 in 8000 for females (Bagni and Zukin, 2019; Garber et al., 2006; Rousseau et al., 2011). Previous high-throughput studies identified only short consensus sites for FMRP binding (ACUK, WGGA, GAC sequences; K=G or U, W=A or U), suggesting that additional elements provide the binding specificity (Suhl et al., 2014). Intriguingly, FMRP was recently identified as a putative m^6^A reader in mammalian cells via unbiased proteomics studies (Arguello et al., 2017; Edupuganti et al., 2017). Furthermore, FMRP-target mRNAs were shown to largely overlap with methylated transcripts (Chang et al., 2017), and FMRP was found to modulate their export to the cytoplasm (Edens et al., 2019; Hsu et al., 2019), as well as their stability (Zhang et al., 2018). These studies indicate roles for FMRP in modulating m^6^A function in the nervous system. However, the physiological relevance of this crosstalk in the context of the FXS has yet to be evaluated. Moreover, it remains to be determined how FMRP interplays with other m^6^A readers.

In the present study, we seek to obtain mechanistic insights into the role of m^6^A in *Drosophila* neurodevelopment. We found that, in addition to controlling axonal growth at neuromuscular junctions, m^6^A prevents axonal crossing and β-lobe fusion of the neurons in the mushroom bodies (MBs), a higher hierarchy circuit of the central brain implicated in a wide range of fly behaviors, including learning and memory. By using an unbiased approach to identify m^6^A readers in the *Drosophila* nervous system, we demonstrate that Ythdf, the unique cytoplasmic YTH protein in *Drosophila*, recognizes methylated transcripts and mediates m^6^A function in restricting axonal growth. We further show that Ythdf directly interacts with Fmr1, the *Drosophila* FMRP homolog, and modulates its binding activity. Ythdf and Fmr1 share common targets related to nervous system development and act in concert to inhibit the translation of positive regulators of axonal growth. Thus, this study demonstrates that Fmr1’s function in axonal growth requires its interaction with the m^6^A reader Ythdf, providing mechanistic insight on this interplay and possibly novel avenues for therapeutic approaches of the FXS.

## Results

### 1- m^6^A restricts axonal growth at the peripheral and central nervous system

Previous studies demonstrated that m^6^A controls several aspects of neuronal development and behavior in *Drosophila melanogaster* (Haussmann et al., 2016; Kan et al., 2017; Lence et al., 2016). In particular, flies lacking Mettl3 are flightless and have reduced speed and orientation defects (Lence et al., 2016). Furthermore, an increased number of synaptic boutons was detected at mutant NMJs. To confirm and extend this initial analysis, we dissected additional alleles of *Mettl3*, as well as *Mettl14* and scored the number of synapses, branches and the overall size of the axons. Consistent with our previous report, we observed an augmentation of synaptic bouton number of about 40% to 50%, depending on *Mettl3* allelic combinations (Fig 1A, B and Supplementary Figure S1A). Furthermore, *Mettl3* mutants displayed significant axonal overgrowth and over-elaboration of synaptic terminals (Fig 1C, D and Supplementary Figure S1B, C). Importantly, all these defects were completely rescued upon expression of *Mettl3* cDNA. Consistent with *Mettl3* loss-of-function phenotypes, the *Mettl14* KO gave identical defects (Fig 1A-D). Thus, these results indicate that m^6^A is required for normal NMJ synaptic architecture in *Drosophila*.

**Figure 1.**
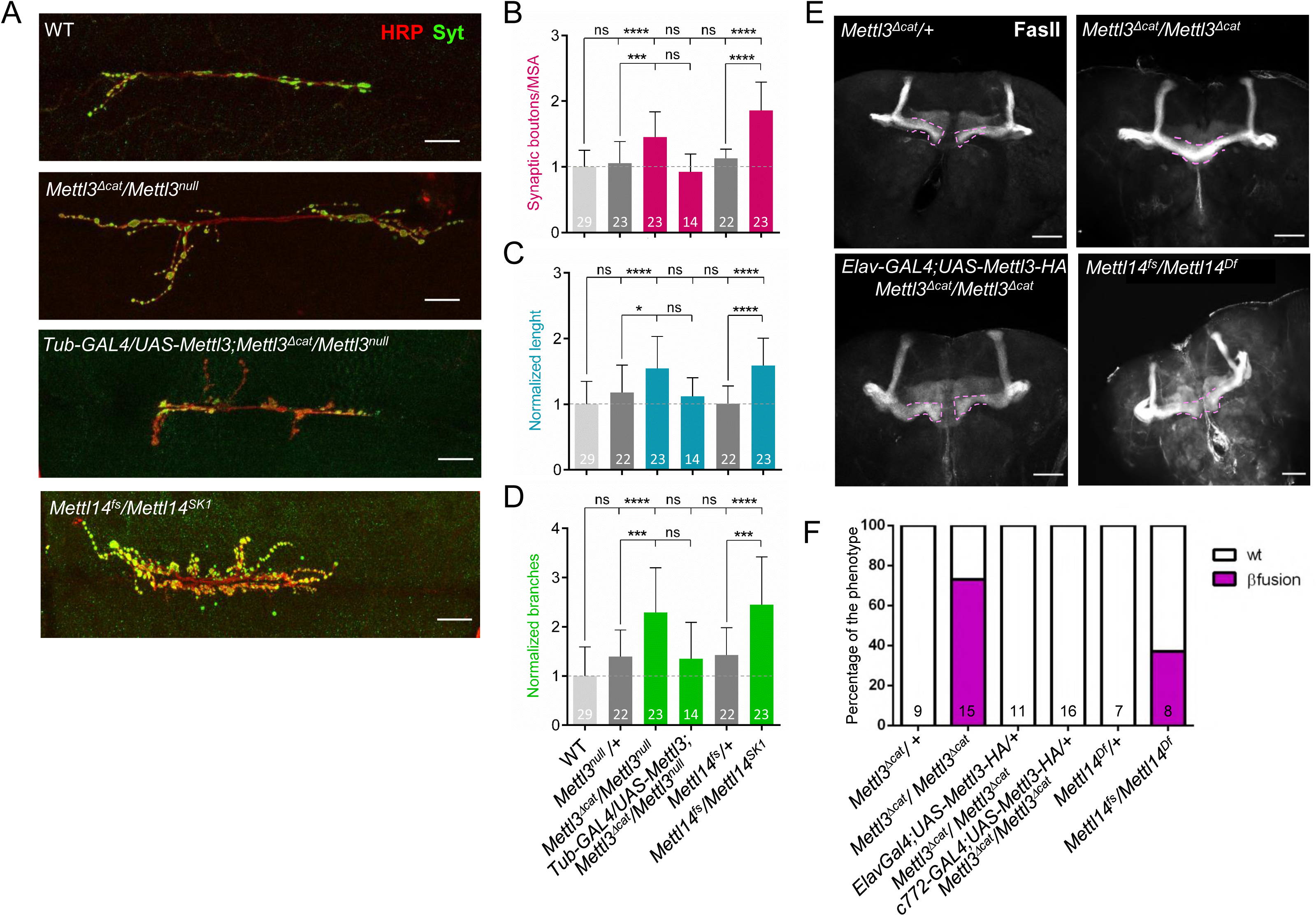
m^6^A regulates axonal growth and guidance in the *Drosophila* nervous system. (A) Representative confocal images of muscle-6/7 NMJ synapses of abdominal hemisegments A2-A3 for the indicated genotypes labelled with anti-Synaptotagmin (green) and HRP (red) to reveal the synaptic vesicles and the neuronal membrane. Scale bar: 20 μm. (B-D) Quantification of normalized bouton number (B, total number of boutons/muscle surface area (μm^2^□×□1,000)), normalized axon length (C) and normalized branching (D) of NMJ 6/7 in A2-A3 of the indicated genotypes. Error bars show mean□±□s.e.m. Multiple comparisons were performed using one-way ANOVA with a post-hoc Sidak-Bonferroni correction. (n.s. = not significant; p<0.05 = *; p<0.01 = **; p<0.001= ***; p<0.0001=****). (E) Immunofluorescence analysis of adult brains for the indicated genotypes using anti-Fascilin II (FasII) antibody to visualize lobes of the MB. Pink dashed lines highlight the normal and fused β-lobes. Scale Bar 50 μM Quantification of the penetrance of fusion phenotype in in the indicated genotypes.

We next asked whether m^6^A was also required for the integrity of the central nervous system (CNS). We dissected adult brains of control and fly mutants for the m^6^A pathway and examined the structure of MBs. Compared to control flies, MBs of *Mettl3* and *Mettl14* KO exhibited midline crossing and fusion of the β lobes (Fig 1E). The penetrance varied from 37% to 73%, depending on the alleles (Fig 1F). A similar defect was observed upon inactivation of *Mettl3* or *Mettl14* specifically in the MB using RNAi (Supplementary Fig S1D, E), suggesting a cell-autonomous requirement of this modification. Furthermore, expression of *Mettl3* cDNA either ubiquitously, pan-neuronally, or in the MBs only, was sufficient to rescue the β lobe overgrowth, confirming the specificity and the cell-autonomous nature of the phenotype (Figure 1E, F). We conclude that m^6^A limits axonal growth in the peripheral and central nervous system.

### 2- Fmr1 and Ythdf bind to methylated sites with different specificity

To decipher the mechanisms underlying the role of the m^6^A pathway in the nervous system we aimed to identify the proteins that mediate m^6^A function in this tissue. We carried out RNA pulldowns in *Drosophila* neuronal cell lysates followed by quantitative mass-spectrometry-based proteomics, as described before (Edupuganti et al., 2017). Briefly, a methylated RNA probe containing four repeats of the m^6^A consensus sequence GGACU was mixed with lysates from BG3 cells, which were derived from the larval CNS. As control, we used the same probe lacking the methylation. After pulldown, bound proteins were subjected to trypsin digestion and analyzed by liquid chromatography - tandem mass spectrometry (LC-MS/MS). Using this approach, we identified eight proteins that were significantly enriched with the methylated probe (Fig 2A). As anticipated, the two YTH domain-containing proteins were among the most strongly enriched. Furthermore, we also found Fmr1, whose mammalian homolog was similarly shown to preferentially bind a methylated probe (Arguello et al., 2017; Edupuganti et al., 2017).

**Figure 2.**
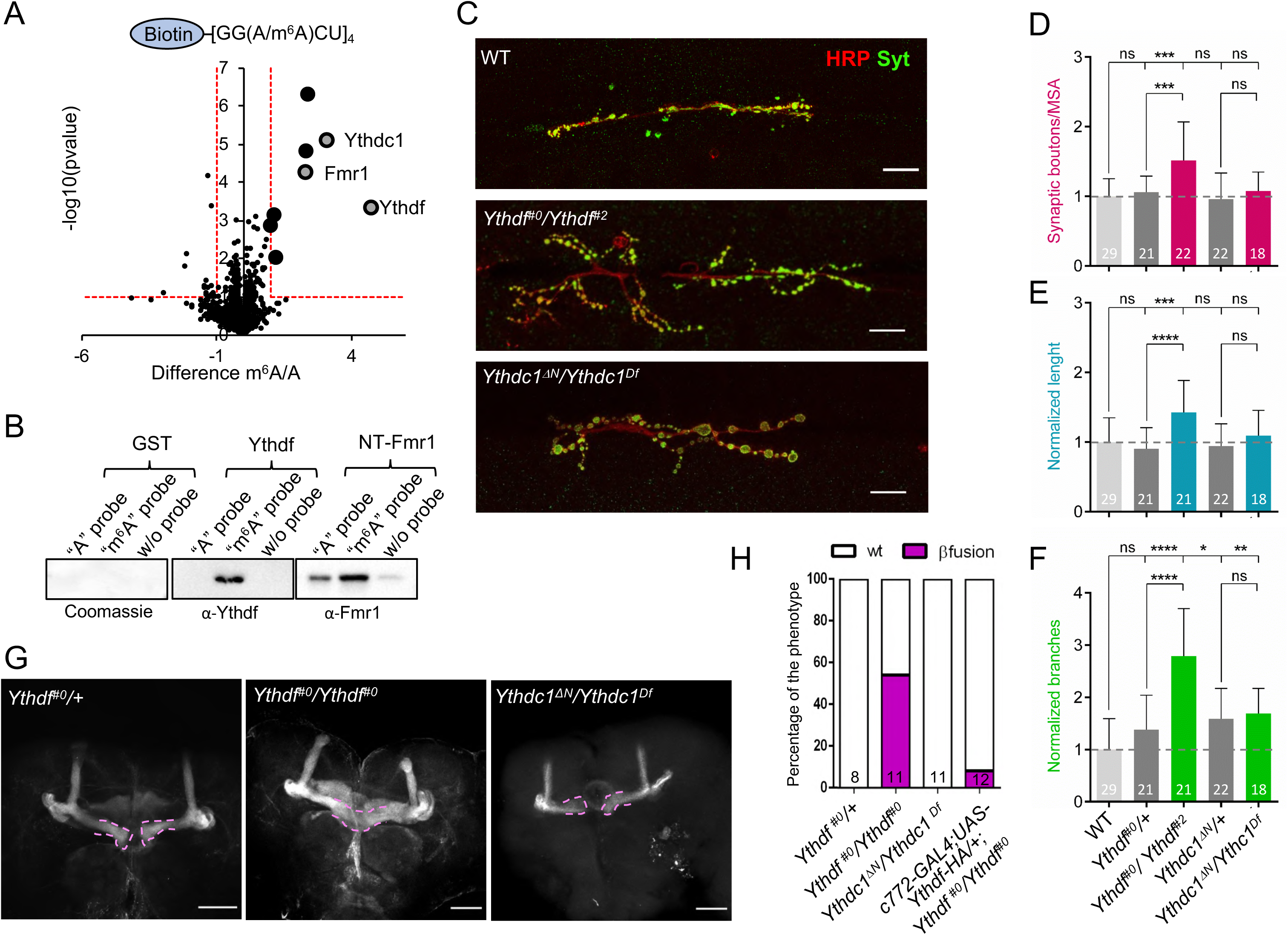
Ythdf and Fmr1 loss of functions are reminiscent of m^6^A loss. (A) Results of m^6^A RNA pulldown in BG3 protein cell extracts. Pulldown were performed using biotinylated probes containing four m^6^A consensus sites (GGACU), with or without the methylation. m^6^A-enriched proteins are depicted by bigger dots. (B) Pulldown using same probes as in (A) and purified recombinant Ythdf and Fmr1 proteins. Both proteins bind more efficiently upon methylation. (C) Representative confocal images of muscle-6/7 NMJ synapses of abdominal hemisegments A2-A3 for the indicated genotypes labelled with anti-Synaptotagmin (green) and HRP (red) to reveal the synaptic vesicles and the neuronal membrane. Scale bar: 20 μm. (D-F) Quantification of normalized bouton number (D, total number of boutons/muscle surface area (μm^2^□×□1,000)), normalized axon length (E) and normalized branching (F) of NMJ 6/7 in A2-A3 of the indicated genotypes. Error bars show mean□±□s.e.m. Multiple comparisons were performed using one-way ANOVA with a post-hoc Sidak-Bonferroni correction (n.s. = not significant; p<0.05 = *; p<0.01 = **; p<0.001= ***; p<0.0001=****). (G) I Immunofluorescence analysis of adult brains for the indicated genotypes using anti-Fascilin II (FasII) antibody to visualize lobes of the MB. For the rescue experiment the MB morphology has been visualized via GFP auto fluorescence driven by the c772Gal4 driver. Pink dashed lines highlight the normal and fused β-lobes. Scale Bar 50 μM. (H) Quantification of the penetrance of fusion phenotype in the indicated genotypes.

Given that binding of Ythdc1 to a methylated probe was already confirmed *in vitro* (Kan et al., 2017), we further aimed to address whether Ythdf and Fmr1 bear the same specificity. Therefore, we purified recombinant GST-tagged Ythdf as well as His-tagged Fmr1 lacking the first 219 N-terminal amino acids (as the full-length version is very unstable, also described in (Chen et al., 2014)), and tested their ability to bind the different probes. Consistent with our pulldown assay from cell lysates, we found that both purified GST-Ythdf and His-Fmr1 bound preferentially to the methylated probe (Fig 2B). While binding of Ythdf was highly specific, a fraction of Fmr1 was also pulled down in the absence of methylation. Interestingly, this behavior was different on RNA probes containing repeats of the alternative m^6^A consensus AAACU. In this case, Fmr1 displayed binding affinity neither for the methylated probe nor for the non-methylated probe, while Ythdf still bound with high affinity the methylated probe (Supplementary Fig S2, Fig 3F). We conclude that the binding of Fmr1 to m^6^A is largely sequence dependent, which is in line with previous pulldown experiments in human cells (Edupuganti et al., 2017) and with a recent study showing that only a fraction of methylated sites is recognized by human FMRP (Hsu et al., 2019).

**Figure 3.**
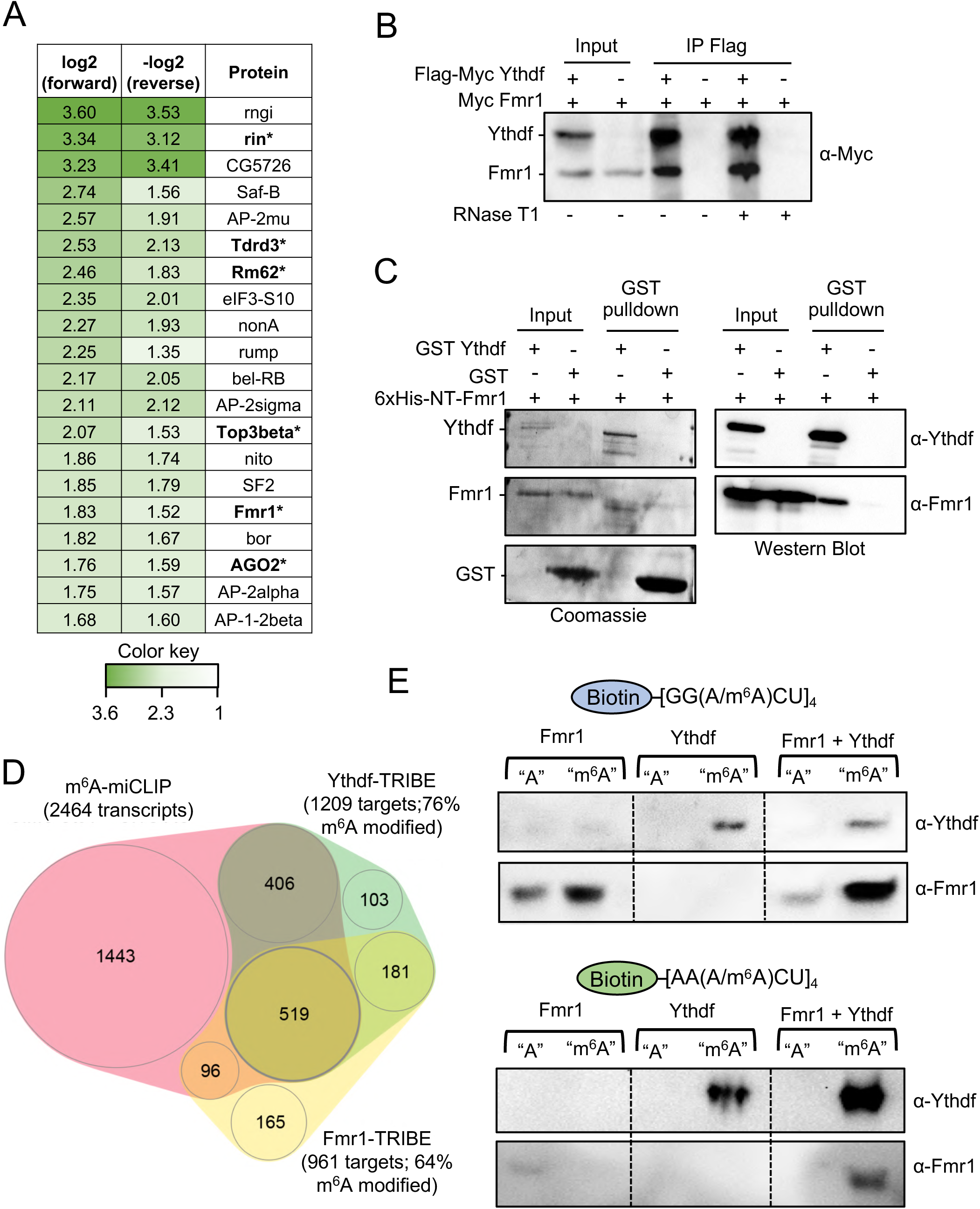
Fmr1 and Ythdf physically interact and share common targets. (A) Heatmap indicating the normalized forward versus inverted reverse experiment enrichments on a log_2_ scale of quantitative proteomics upon pulldown of Flag-tagged Ythdf in S2R+ cells. The threshold was set to a 1-fold enrichment (dashed line). Fmr1 is co-purified with Ythdf. Known Fmr1 interacting proteins are highlighted in bold (B) Co-immunoprecipitation experiment in S2R+ cells co-expressing FlagMyc-tagged Ythdf and Myc-tagged Fmr1. Ythdf was used as a bait via its Flag tag. The lysate was treated with RNase T1 as indicated to remove interactions enabled by RNA. (C) GST pulldown of recombinant GST-Ythdf or GST alone mixed with recombinant 6xHis-NT-Fmr1. The purified proteins were analyzed by coomassie staining and immunoblotting using α-Ythdf and α-Fmr1 antibodies. (D) Overlap of miCLIP dataset generated in S2R+ cells defining m^6^A modified transcripts with Fmr1 and Ythdf mRNA targets determined by TRIBE in S2R+ cells. (E-F) Pulldown of biotinylated RNA probes of repetitive GGACU sequences (E) or AAACU sequences (F) incubated with *E. coli* lysate expressing recombinant GST-Ythdf or/and 6xHis-NT-Fmr1. Pulled down proteins were analyzed by immunoblotting using α-Ythdf and α-Fmr1 antibodies.

### 3- Fmr1 and Ythdf limit axonal growth

The *Fmr1* loss of function was previously shown to give overgrowth at NMJs, as well as fusion of MB β lobes (Supplementary Fig S3 and (Michel et al., 2004; Zhang et al., 2001)), suggesting that it could mediate the m^6^A axonal growth function. To address whether Ythdc1 and/or Ythdf also control NMJ morphology, we dissected third instar larvae carrying mutations in the *Yth* genes. Using our previously described *Ythdc1* allele combined over a deficiency line spanning the locus we did not detect any gross morphological defect (Fig 2C-F). To address the contribution of Ythdf, we generated mutant alleles using the CRISPR/Cas9 approach (Supplementary Fig S4, see also material and methods section). Examination of the NMJs in the trans-heterozygote flies revealed significant overgrowth compared to control flies (Fig 2C-F). Thus, these results indicate that in addition to Fmr1, Ythdf may also contribute to the m^6^A-dependent regulation of NMJ morphology.

To address the role of YTH proteins in the CNS, we dissected adult brains of control and fly mutants for the respective *Yth* genes and examined the MB structure. Compared to control flies, *Ythdf* KO brains show a substantial fusion of the β lobes (55%), mimicking the loss of *Mettl3* and *Mettl14* (Fig 2G, H). This defect was rescued upon Ythdf overexpression. In contrast, the lack of Ythdc1 displayed no visible defect. Altogether, these results indicate that the m^6^A pathway controls axonal growth, both at NMJs and MBs, possibly via Fmr1 and Ythdf.

### 4- Ythdf interacts physically with Fmr1

To address how Ythdf prevents overgrowth at NMJs and MBs, we searched for co-factors using stable isotope dimethyl labeling-based quantitative proteomics upon immunoprecipitation of Flag-tagged Ythdf from S2R+ cells, which is an embryonic derived cell line. We identified 51 factors that showed more than two-fold enrichment in the Flag-Ythdf pulldown fraction in comparison to a control pulldown (Supplementary Fig S5A). The co-purified proteins were especially enriched for RNA binding proteins and translation regulators (Supplementary Fig S5B). Interestingly, Fmr1 was amongst the 20 most enriched proteins (Fig 3A). In fact, several proteins previously shown to interact with Fmr1 were also pulled down with Flag-Ythdf (Fig 3A, highlighted in bold; (Ishizuka et al., 2002; Sahoo et al., 2018; Xu et al., 2013)), suggesting that Ythdf may be part of a whole Fmr1 complex.

To validate the co-existence of Ythdf and Fmr1 in the same complex, we performed co-immunoprecipitation assays in S2R+ cells ectopically expressing Flag-Myc-tagged Ythdf and Myc-tagged Fmr1 in the presence of RNase. Notably, these experiments confirmed that Fmr1 and Ythdf co-precipitate, independently of RNA (Fig 3B). To next address if these two proteins interact directly, we tested whether purified recombinant GST-tagged Ythdf and Fmr1 proteins could pull down each other. As shown in Fig 3C, Fmr1 could be co-purified by pulling down GST-Ythdf but not GST alone, indicating that Ythdf and Fmr1 directly interact.

### 5- Ythdf and Fmr1 regulates translation of similar targets

In order to identify Ythdf and Fmr1 mRNA targets, we used the TRIBE approach, as previously described (Worpenberg et al., 2019). Briefly, metal-inducible fusion constructs expressing Ythdf-cdAdar, Fmr1-cdAdar or cdAdar alone were transfected in S2R+ cells and RNA was isolated for sequencing two days after the induction of the constructs. Bound mRNA targets were identified by scoring A-I editing events obtained after comparison with unspecific events generated by cdAdar alone. Using a stringent cut-off, we identified 1209 Ythdf and 961 Fmr1 targets, with a significant degree of overlap (n=645, Fig 3D, Supplementary Table 1). To address whether these targets were methylated, we performed miCLIP-seq using S2R+ cell extracts. We identified 2464 methylated transcripts (about 34% of expressed genes), with strong enrichment of m^6^A sites within 5’ UTR, in the sequence context AAACA (Supplementary Fig S6A, B; Supplementary Table 2). This profile is distinct from vertebrate and is consistent with an earlier report (Kan et al., 2017). Among methylated transcripts, 925 were common with Ythdf (76% of Ythdf targets) and 615 with Fmr1 (64% of Fmr1 targets). Even though S2R+ cells do not have a neural origin, common methylated targets were enriched for axon regeneration (Supplementary Fig S6C). Moreover, consistent with the *in vivo* phenotypes, genes enriched for regulation of microtubule depolymerization were overrepresented. Altogether, these experiments identify common methylated targets for Ythdf and Fmr1, suggesting they could act together to regulate gene expression.

To further understand the interplay between Ythdf and Fmr1 on RNA, we repeated pulldown experiments using the aforementioned biotinylated RNA probes. Combining GST-Ythdf and His-Fmr1 did not substantially alter their respective binding affinity to RNA probes in the GGACU sequence context, independently of the methylation status (Fig 3E). However, adding GST-Ythdf to His-Fmr1 enabled Fmr1 binding to the methylated AAACU RNA probe (Fig 3F). Binding of Ythdf also appeared increased. These results indicate that Ythdf can facilitate binding of Fmr1 to a subset of methylated sites that diverges from the preferred Fmr1 binding site GGACU. Collectively, these experiments led to the identification of common methylated targets of Ythdf and Fmr1, and strongly suggest that these two factors directly interact to regulate gene expression.

### 6- Ythdf and Fmr1 control axonal growth at the MB by inhibiting *chic* translation

Among the common targets of Fmr1 and Ythdf that present several methylation sites were *chickadee* (*chic*) transcripts. *chic* codes for Profilin, an actin-binding protein that modulates many processes depending on actin dynamics, among these neuronal growth and remodeling (Verheyen and Cooley, 1994). Importantly, previous work by Reeve and colleague demonstrated that Fmr1 controls axonal growth in the CNS via inhibition of *chic* translation (Reeve et al., 2005). We first performed an RNA immunoprecipitation assay (RIP) to confirm the binding of Fmr1 and Ythdf to *chic* RNA in the adult brain using HA-tagged Ythdf and GFP-tagged Fmr1 driven by the neuronal driver *elav*-GAL4. As shown in Supplementary Fig 7A, both proteins were able to pulldown *chic* RNA. To test if the m^6^A pathway regulates *chic* expression in a similar manner as Fmr1, we performed immunostaining of brains from WT and *Ythdf* mutant. We found that the lack of Ythdf led to increased Profilin levels (Fig 4A, B). This was also confirmed by Western blot analysis (Fig 4C, D). In contrast, RNA levels remained unaffected (Fig 4E). These results prompted us to investigate if the increased Profilin level was responsible for the β-lobe fusion phenotype. We then crossed *Ythdf* mutant to *chic* mutant flies and scored for MB developmental defects. As shown in Fig 4F and G, loss of one copy of *chic* was sufficient to rescue the β-lobe fusion phenotype of *Ythdf* mutant flies. On the other hand, *chic* heterozygote flies already showed β-lobe fusion, suggesting that the level of Profilin has to be tightly regulated to ensure proper brain wiring. To dissect the molecular mechanisms underlying the increased Profilin level we performed RIP assays by immunoprecipitating endogenous Fmr1 from WT and *Ythdf* mutant brains. We observed a significant reduction in the *chic* RNA immunoprecipitated by Fmr1 in the absence of Ythdf (Fig 4H). Hence, this result suggests that Ythdf facilitates binding of Fmr1 to *chic* RNA, thereby inhibiting its translation.

**Figure 4.**
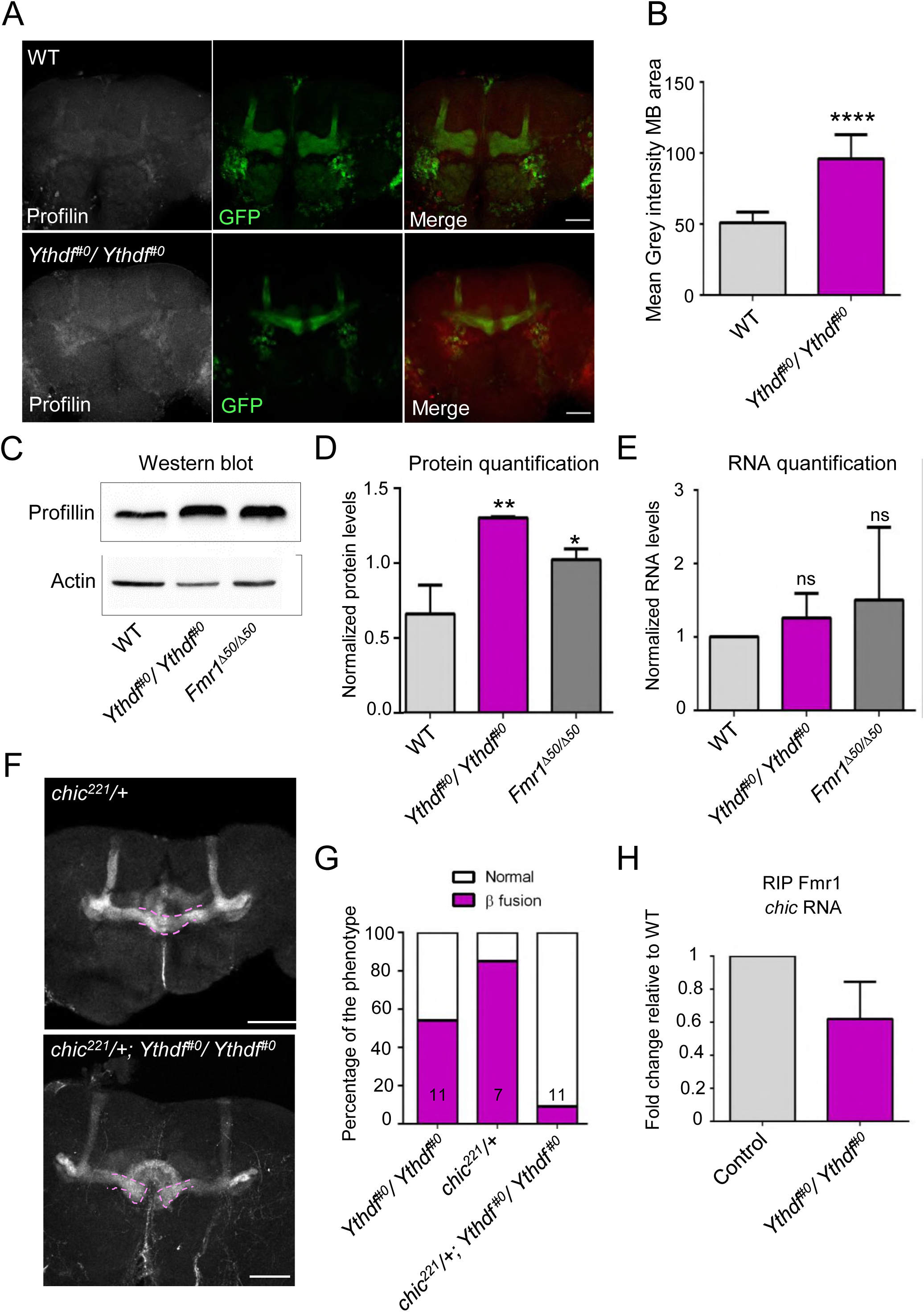
Fmr1 and Ythdf regulate CNS axonal growth by modulating Profilin levels. (A) Immunofluorescence analysis of control brains *c772Gal:UAS-CD8-GFP* and *Ythdf* mutant *c772Gal;UAS-CD8-GFP;Ythdf*^Δ*YTH*^ with anti-Profilin (Grey/Red) and anti-GFP (Green) antibodies. Scale bar: 50 μM (B) Mean grey intensity in the MB area measured with Fiji using GFP to define the region of interest. Average of 6-13 brains per genotype. (C) Representative Western blot analysis of protein extracts of late pupae heads (85-95h) from control, *Ythdf^#0^/ Ythdf^#0^* and *Fmr1*^Δ*50*^/*Fmr1*^Δ*50*^ flies. The membranes were probed with anti-Profilin or anti-Actin antibodies. (D) Quantification of Profilin protein levels obtained from three independent protein-extraction and Western blot analysis as in C. Quantification was performed using Fiji. p<0.005 = **; p<0.05 = *, measured with unpaired t-test. (E) Quantification of *chic* mRNA levels obtained from six independent RNA extractions from control, *Ythdf*^Δ^*^YTH^* and *Fmr1*^Δ*50*^ late pupae heads, via real-time PCR analysis. Statistical analysis using unpaired t-test show no significant difference between the samples. (F) Immunofluorescence analysis of adult *chic/+* and *chic/+; Ythdf^#0^/Ythdf^#0^* using anti-FasII antibody. Scale Bar: 50 μM (G) Quantification of the penetrance of β-lobe fusion phenotype for the indicated genotypes H) RNA immunoprecipitation assay. Quantification of *chic* mRNA levels upon immunoprecipitation of endogenous Fmr1 in control and *Ythdf^#0^/Ythdf^#0^* late pupae heads (100 heads per genotype) with Real-time PCR. The graph represents the average of three independent biological repeats. p <0.05 = *, measured with unpaired t-test.

### 7- Ythdf and Fmr1 control axonal growth at the NMJ by inhibiting *futsch* mRNA translation

We next investigated whether *chic* could also be the common effector of Fmr1 and Ythdf at the NMJ. As in the adult brain, we found that both proteins interact with *chic* RNA in the larval nervous system (Supplementary Fig S7B). However, in contrast to what we observed at the MB, removing one copy of *chic* had no effect on the NMJ phenotype produced by the loss of Ythdf (Supplementary Fig 7C). These results indicate that *chic* is not responsible for the m^6^A-dependent phenotype at the NMJ, and that other target(s) must be involved. To reveal their identity, we generated transgenic flies expressing Ythdf-cdAdar under the control of an UAS promoter. The construct was specifically expressed in the nervous system using the *elav*-GAL4 driver and, after dissection, RNA was isolated and submitted for high-throughput sequencing. Using this strategy, we identified 982 Ythdf target mRNAs (Supplementary Table 3). Interestingly, among them was *futsch*, which encodes a microtubule associated protein that is a key target of Fmr1 in the regulation of axonal growth at the NMJ (Zhang et al., 2001). We confirmed the binding of Ythdf and Fmr1 to *futsch* mRNA in larval brains by immunoprecipitating HA-tagged Ythdf and GFP-tagged Fmr1 driven by the neuronal driver *elav*-GAL4 (Supplementary Fig S7B). Remarkably, in *Mettl3* and *Ythdf* mutants, Futsch protein level was significantly upregulated at the larval NMJ and in larval brain extracts, which is reminiscent of the previously described upregulation observed in the *Fmr1* mutant (Fig 5A-C). In contrast, *futsch* mRNA level was downregulated by two-fold (Fig 5D). To discriminate between a defect in translation inhibition and a defect in protein decay, we performed Translating Ribosome Affinity Purification (TRAP). By immunoprecipitation of a GFP-ribosomal fusion protein, TRAP enables the isolation of mRNAs associated with at least one 80S ribosome, providing an estimation of the translation status of associated transcripts. We expressed RPL10-GFP in neurons using the *elav*-GAL4 driver and pulled down associated RNA using an anti-GFP antibody. Using this approach, we found that *futsch* mRNA was strongly enriched in the *Ythdf* mutant, demonstrating that its translation was likely increased (Fig 5E). Altogether, these experiments indicate that, like Fmr1, the m^6^A pathway restricts *futsch* mRNA translation in the larval nervous system.

**Figure 5.**
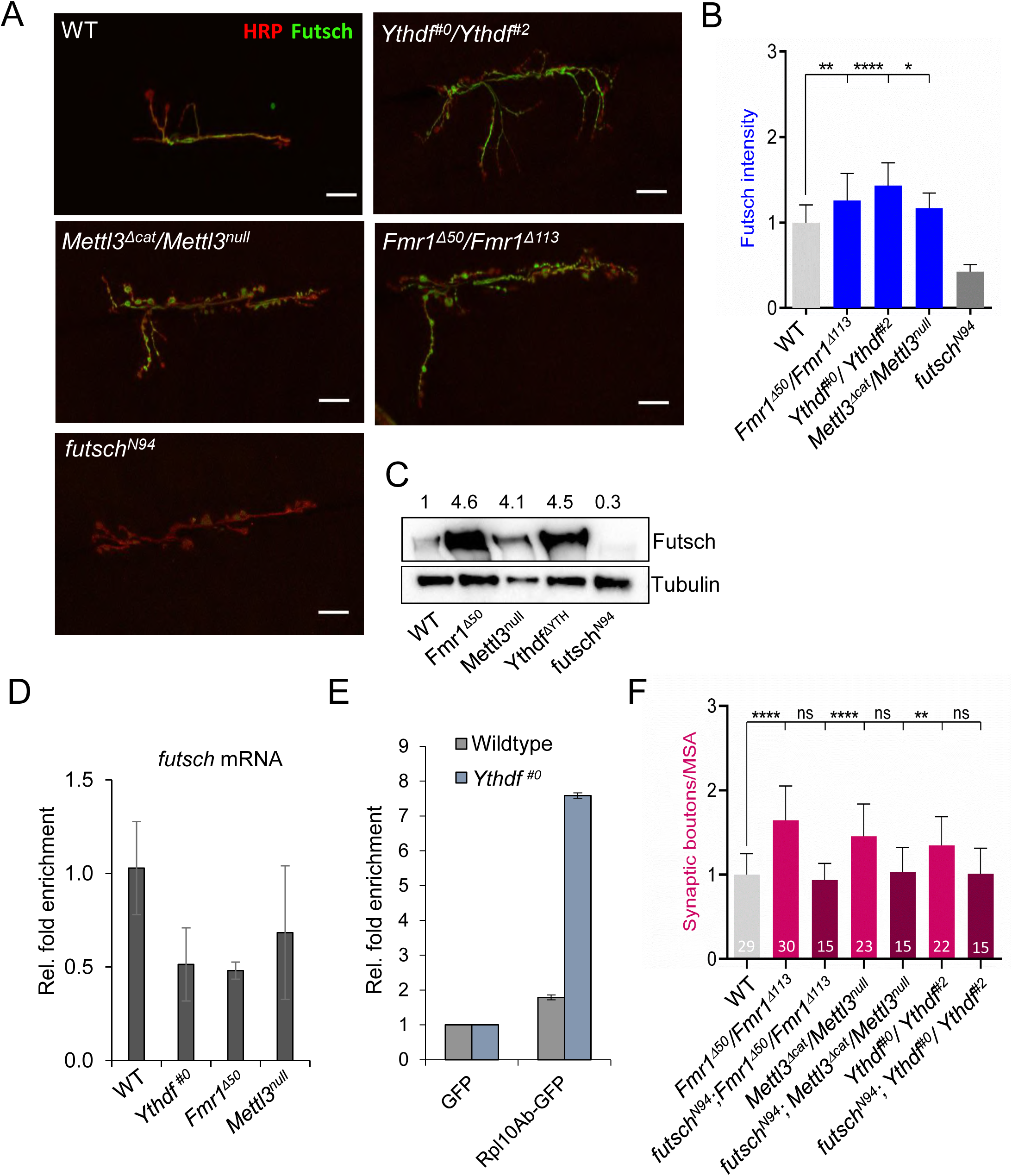
Fmr1 and Ythdf regulate CNS axonal growth by modulating Futsch levels. (A) Representative confocal images of muscle-6/7 NMJ synapses of abdominal hemisegment A2 for the indicated genotypes labelled HRP (red) and with α-futsch (green) to reveal the neuronal membrane and Futsch protein level. (B) Quantification of normalized Futsch protein level at NMJ 6/7 in A3 of the indicated genotypes. Error bars show mean□±□s.e.m. *P* values were determined with a Student’s *t*-test. (n.s. = not significant; p<0.05 = *; p<0.01 = **; p<0.001= ***; p<0.0001=****). (C) Western Blot analysis of Futsch protein level in isolated third instar larval brains of different mutants compared to control. Numbers indicate normalized Futsch protein level in comparison to Tubulin levels. (D) Quantification of *futsch* mRNA levels obtained from isolated 3^rd^ instar larvae brains of the indicated genotypes via real-time PCR analysis. (E) Translating Ribosome affinity purification assay. Quantification of *futsch* mRNA levels upon immunoprecipitation of GFP-tagged Rpl10Ab or GFP in wildtype and *Ythdf^#0^* 3^rd^ instar larvae with Real-time qPCR. (F) Quantification of normalized bouton number (total number of boutons/muscle surface area (μm^2^□×□1,000)) of NMJ 6/7 in hemisegment A3 of the indicated genotypes.

To functionally address the relationship between the m^6^A pathway and *futsch* mRNA at the NMJ, we performed genetic experiments. As shown earlier, the number of synapses in the *Fmr1* mutant was restored to a normal level by removing *futsch* function (Fig 5F, (Zhang et al., 2001)). Similar rescues were obtained in the *Mettl3* and *Ythdf* mutants (Fig 5F). This indicates that the m^6^A pathway controls NMJ morphology, primarily via repressing *futsch* mRNA translation.

### 8- Ythdf binding to *futsch* mRNA 5′ UTR promotes translation inhibition

To functionally investigate the mechanism underpinning the influence of m^6^A and of the Ythdf/Fmr1 complex on *futsch* translation, we determined the location of the m^6^A sites on the *futsch* transcript. By analyzing our miCLIP-seq data, we found that *futsch* harbors two m^6^A sites in its 5′ UTR, close to the start codon (Fig 6A). We verified the methylation of one of these sites *in vivo* by performing single-base elongation and ligation-based qPCR amplification (SELECT) (Xiao et al., 2018) using larval brain extract (Supplementary Fig 7D). To next define the influence of the methylation on *futsch* expression, we designed reporters containing the GFP coding sequence downstream of the WT *futsch* 5′ UTR or a version containing A-T point mutations at the m^6^A sites (Fig 6B). We tested whether the binding of Ythdf and Fmr1 to the mutated reporter transcript was altered by immunoprecipitating Flag-tagged Ythdf and Fmr1 from S2R+ cell lysate co-transfected with the reporter constructs. A significantly lower amount of the mutated reporter transcript was recovered in the pulldown (Fig 6C). In addition, we found that the enrichment of the *futsch* 5′ UTR reporter in the Fmr1 pulldown fraction was significantly increased when Ythdf was co-expressed (Fig. 6D), indicating that Ythdf increases the binding of Fmr1 to its targets, likely by direct recruitment. We next analyzed if mutations of the two m^6^A sites led to a change in translation of the reporter, similar to the observed increased translation of endogenous *futsch* in *Ythdf* mutants. As shown in Fig 6E, F, the GFP level expressed from the mutated reporter was increased by two-fold in comparison to the WT reporter. Therefore, we conclude that the *futsch* 5′ UTR harbors at least two m^6^A sites, which are required for Ythdf and Fmr1 binding and for the translational repression of *futsch* (Fig 6G).

**Figure 6.**
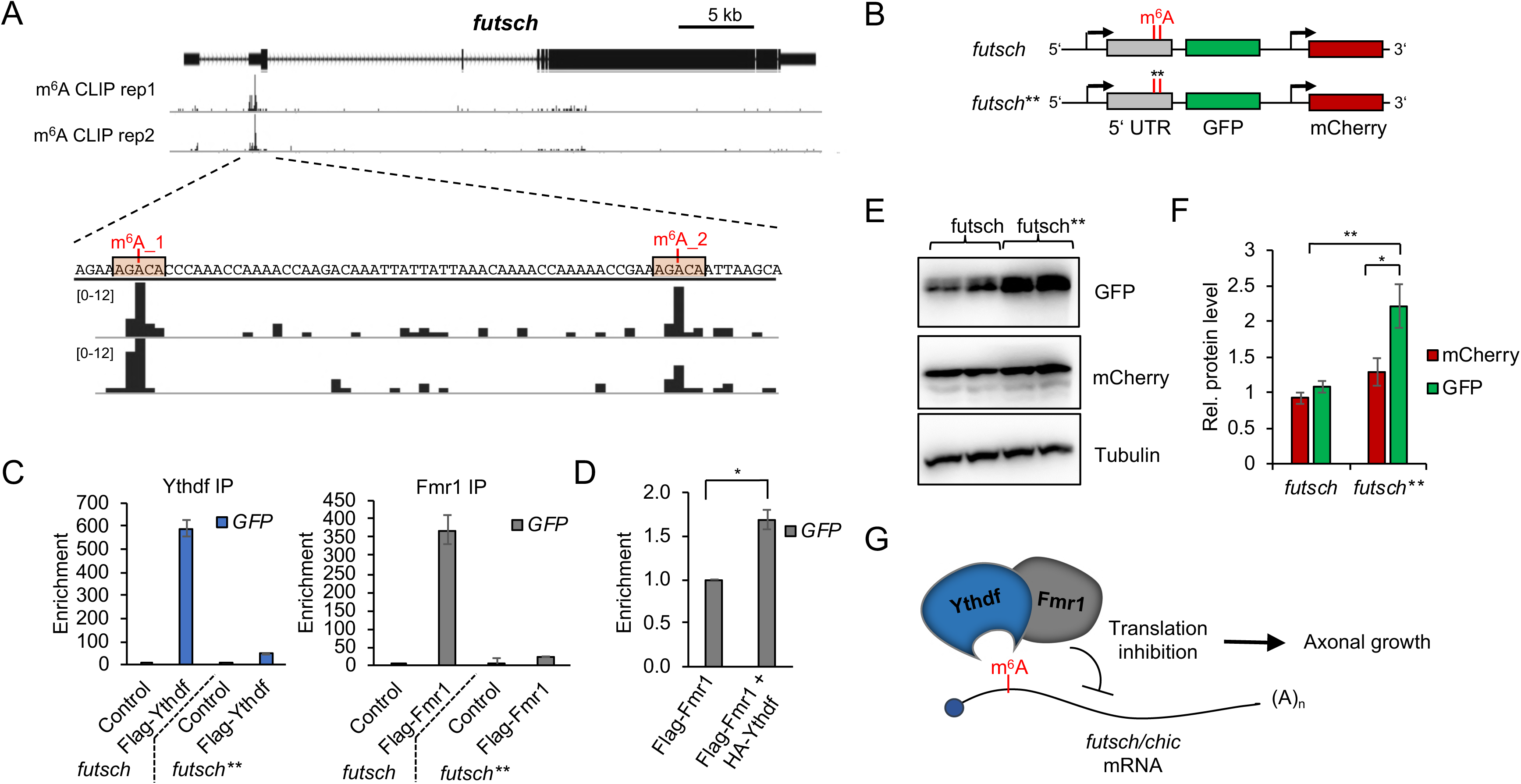
Fmr1 recruits Ythdf to repress Futsch translation. (A) *futsch* genomic region (X1,408,435 – 1,454,00) and corresponding browser tracks from miCLIP-seq experiments in S2R+ cells. *futsch* transcripts contain two m^6^A peaks in its 5′ UTR, as indicated. (B) Schematic of GFP reporter constructs containing an *actin* promoter and either wildtype 5′ UTR region of *futsch* or a mutated version (*futsch*\*\*) containing two-point mutations (A – T) at the identified m^6^A sites. mCherry is under the control of a same *actin* promoter. (C, D) RNA immunoprecipitation assay. Quantification of *GFP* RNA levels upon immunoprecipitation of Flag-tagged Fmr1 or Flag-tagged Ythdf (C) or Flag-tagged Fmr1 in the presence of overexpressed HA-tagged Ythdf (D). *mCherry* levels were used as a normalization control. (E, F) Representative Western blot analysis (E) and quantification (F) of protein levels from S2R+ cells transfected with the wildtype or mutated *futsch* reporter constructs. (G) Model of the interplay between Ythdf and Fmr1 in translation control.

## Discussion

m^6^A on mRNA is emerging as a key modulator of nervous system biology (for recent review see (Livneh et al., 2020)). Despite the increasing amount of data associating m^6^A function to brain development, neuronal differentiation, regeneration and synaptic function, the molecular mechanisms underlying these functions remain largely incomplete. Here we show that m^6^A is required for proper neuronal development in both CNS and PNS, where it prevents MB β-lobes fusion and NMJ overgrowth, respectively. We demonstrate that m^6^A exerts these functions via a critical interplay between Ythdf and Fmr1. We show that Ythdf recruits Fmr1 to a subset of mRNA to inhibit their translation and ensures proper axonal growth and homeostasis. Hence, our study reports that m^6^A controls the development of the nervous system by modulating Fmr1 target selectivity.

### m^6^A in axon growth and guidance

Previous work showed that flies lacking m^6^A are flightless and have reduced locomotion, due to impaired neuronal functions (Haussmann et al., 2016; Kan et al., 2017; Lence et al., 2016). In this study, we found that m^6^A controls axonal growth and guidance in the PNS and CNS, by regulating protein levels of two key components of the cytoskeleton, Futsch and Profilin, respectively. Roles for m^6^A in axonal growth and guidance have been previously observed in mammals. For instance, depletion of FTO in axons of dorsal root ganglia neurons represses axon elongation in mouse (Yu et al., 2018). In this model, FTO depletion inhibits axon growth due to increased m^6^A levels on *growth-associated protein-43 (GAP-43)* mRNA, resulting in reduced GAP-43 protein abundance. Accordingly, the growth defect can be rescued by expressing a deficient-m^6^A construct encoding GAP-43. However, how m^6^A inhibits *GAP-43* translation is not understood. Perhaps a similar mechanism as described in our work, involving the recruitment of FMRP, is operating (see also below). This would be consistent with the upregulation of GAP-43 observed at synapses of *Fmr1* KO mice (Klemmer et al., 2011). Upon injury of the same dorsal root ganglion neurons, it was shown that m^6^A levels increase dramatically, resulting in enhanced protein synthesis and eventually axon regeneration (Weng et al., 2018). A similar activity was also observed in the adult CNS. Thus, m^6^A plays a positive role in axonal growth upon injury, which seems in contradiction to its role in normal growth during development. Nevertheless in mouse embryonic dorsal spinal cord (DSC), m^6^A is required for axon growth and guidance by promoting *Robo3.1* translation via YTHDF1 (Zhuang et al., 2019). Thus m^6^A has the ability to either promote or repress axon growth and guidance, which depends on the developmental and physiological contexts.

### Ythdf controls axon growth and guidance through Fmr1

In mammals, there are three YTHDF proteins that were shown to control translation and mRNA decay. The current view is that YTHDF1 stimulates translation, YTHDF2 decreases mRNA stability while YTHDF3 regulates both processes (Patil et al., 2018; Zhao et al., 2019). Nevertheless, this is probably an over-simplification, as for instance, YTHDF1 depletion can also result in mRNA decay (Wang et al., 2015). Thus, more work is required to appreciate how the three YTH proteins interplay in the cytoplasm and how much of their functions overlap. In *Drosophila*, there is only one Ythdf protein but its function was not characterized prior to our work. In fact, the role of cytoplasmic m^6^A in this organism has remained enigmatic. Our attempt to demonstrate a role on RNA stability in S2R+ cells was not conclusive, so it is unclear whether m^6^A regulates this process in flies (data not shown). The fact that most m^6^A sites resides in 5′ UTR, near the start codon, suggests instead that the main role of cytoplasmic m^6^A in *Drosophila* is to regulate translation. Consistent with this assumption, we found that m^6^A controls *futsch* and *chic* translation through Ythdf activity. However, in contrast to mammalian YTHDFs, *Drosophila* Ythdf does not activate their translation. It represses translation of these mRNA via recruitment of the translation inhibitor Fmr1. For future studies, it would be interesting to address whether translation inhibition is the major function of Ythdf in *Drosophila*, or whether it has a broader role depending for instance on the nature of its interacting partners. Furthermore, whether such inhibitory role of YTHDFs in translation also exists in mammals awaits future investigations.

FMRP was recently shown to bind methylated RNA and to facilitate their export (Edens et al., 2019; Hsu et al., 2019), as well as to protect them from degradation by preventing YTHDF2 binding (Zhang et al., 2018). It was proposed to be an m^6^A reader, acting in a sequence dependent context. Indeed, *in vitro* pull down as well as FMR1 CLIP demonstrated association to GGACA/C/U sequence, with clear enrichment upon methylation (Arguello et al., 2017; Edupuganti et al., 2017; Hsu et al., 2019). Whether this enrichment is due to direct recognition of the methyl group by FMR1 or to a change in RNA accessibility is still unclear. Replacing the first or the second G by A strongly decreased FMR1 association, while no effect was observed on YTH binding. Here we show that similar rules apply in *Drosophila*. In addition, our data further demonstrate that Ythdf interacts directly with Fmr1 and stabilizes its interaction to RNA, especially when sites diverge from the GGACU sequence context. This direct interaction is essential for both protein activities in the context of axonal growth. Of note, human YTHDF2 was recently shown to be in a same complex with FMR1, suggesting that this mode of interaction is conserved (Zhang et al., 2018). Nevertheless, the fact that the AAACU context is more prominent in *Drosophila* suggests that Fmr1 dependency towards Ythdf may even be more critical in this organism. To verify this prediction, it would be necessary to address Fmr1 binding on a global scale in the absence of m^6^A or Ythdf.

### Relevance of this interplay in FXS

The absence of FMRP leads to the FXS, which is a severe inherited neuronal disorder that currently lacks efficient therapeutic treatment. The phenotype of the patients suffering from FXS is often complex, accompanied by an increase in autism spectrum disorder specific traits and other features like delayed motor development, hyperactivity, aggression and epileptic seizures (reviewed in (Dahlhaus, 2018; Garber et al., 2008; Hagerman et al., 2014; Hagerman et al., 2002; Kidd et al., 2014; Maurin et al., 2014; Santoro et al., 2012; Schaefer et al., 2015; Utari et al., 2010)). These abnormalities result from defects in neuronal development and maturation. Interestingly, the phenotypes of *Fmr1* mutants are reminiscent of the pathogical symptoms of FXS patients and, consequently, *Drosophila* has been widely used to learn the basic mechanisms underlying FMRP functions and to test the efficacy of drug treatment (Drozd et al., 2018). In particular, the increased synapse arborization and bouton number at the NMJ recall the dendritic spine overgrowth observed in FXS patients. Morevover, expressing human FMR1 in *Drosophila* Fmr1 null mutants rescues the overgrowth at the NMJ and the defect in the brain, highlighting the functional conservation of the two orthologues (Coffee et al., 2010). Since FMR1 is involved in different functions such as splicing, nuclear export and translation, it remained unclear which activity was more relevant in the FXS etiology. A clue came when treatment with the translation inhibitor puromycin could rescue several aspects of FXS, including the locomotion phenotype and the overgrowth at the NMJ, suggesting that it was an excess of translation that yields these defects (Bolduc et al., 2008; Kashima et al., 2017; Stefani et al., 2004). Accordingly, the first target identified for Fmr1 was Futsch, a microtubule-associated protein orthologue of mammalian MAP1B (Zhang et al., 2001). Fmr1 negatively regulates *futsch* translation and this is necessary to prevent NMJ overgrowth. Importantly, this function was also found in mice (Lu et al., 2004). Our data showing that Fmr1 represses *futsch* translation via m^6^A activity directly links m^6^A to some aspects of the FXS studied in the fly model. It would be of critical importance to test whether a similar mechanism also applies to mammals. The FMR1-mediated nuclear export and stability of methylated RNA as recently uncovered may contribute as well to the disease (Edens et al., 2019; Hsu et al., 2019; Zhang et al., 2018). It is worth mentioning that while our study mainly focuses on two key Fmr1 targets involved in the gross morphology of the nervous system, it is likey that m^6^A and Fmr1 regulate additionnal targets involved in more subtle processes such as synapse functionality and complex trait behaviors.

In conclusion, our study indicates that m^6^A modulates both CNS and PNS development by restricting axonal growth and promoting correct assembly of the neural circuits. These functions are reminiscent to the functions of Fmr1 in the nervous system and our work show that both m^6^A and Fmr1 tightly cooperate to regulate these processes. We foresee that this new knowledge will open new avenues for the design of complementary treatments of FXS.

## Materials and Methods

### *Drosophila* stocks

The stocks used are the following: *w1118, Fmr1^Delta50M^*, *Fmr1^Delta113M^, chic^221^, futsch^N94^*, *Mettl14^Df^, Ythdc1^Df^, Tub-GAL4, Elav-Gal4, c772Gal4:UAS-CD8-GFP, Elav-Gal4;;UAS-FMR1-GFP/TM3* (Bloomington Stock Collection), *Mettl3^null^*, *Mettl3^Deltacat^*, UAS-Mettl3, *Mettl14^fs^*, *Ythdc1^DeltaN^* (Lence et al., 2016)*, Mettl3^SK2^*, *Mettl14^SK1^* (kind gift from Eric Lai), *P(GD9882)v20969/TM3* (VDRC), *P(GD9882)v20968/TM3* (VDRC), w1118;P(GD11887)v27577 (VDRC), w1118;P(GD16300)v48560 (VDRC).

*Drosophila melanogaster* Canton-S with mutant alleles for *Ythdf* were generated using the CRISPR/Cas9 system, as described previously (Lence et al., 2016). Guide RNA sequences used were CTTCGGATAAATTCTTTCCGAATA and AAACTATTCGGAAAGAATTTATCC as well as CTTCGGGCGAGTGGGGCAGGCGCG and AAACCGCGCCTGCCCCACTCGCCC. The first allele (*Ythdf^#0^*) produces a deletion of 1221 nucleotides in the coding sequence, deleting residues 172 to 557, that includes the whole YTH domain. The second allele (*Ythdf^#2^*) is a deletion of 1319 nucleotides and removes residues 162 to 558.

### *Drosophila* cell lines

*Drosophila* S2R+ are embryonically derived cells obtained from the *Drosophila* Genomics Resource Center (DGRC; Flybase accession FBtc0000150) while *Drosophila* BG3 cells are derived from central nervous system of third instar larvae (DGRC; Flybase accession FBtc0000068). Mycoplasma contamination was not detected (verified by analyzing RNA sequencing data).

### Immunohistochemistry

For the immunohistochemistry in adult, pupal and larval brains the following protocol was used: brains were collected in cold PBS and subsequently fixed for 10 minutes in 4% Formaldehyde (in PBS 0.3% Triton X-100). Upon three washes in PBS 0.3% Triton X-100 the brains were blocked in 10% BSA (in PBS 0.3% Triton X-100) for 1 h rocking at RT. After this step, the brains were incubated overnight at 4°C with the primary antibody appropriately diluted in blocking solution. The second day the samples were washed 3 times in PBS 0.3% Triton X-100 and then incubated with the appropriate secondary antibody diluted in blocking solution for 1h at RT. Upon three washes in PBS 0.3% Triton X-100 the brains were mounted and kept at 4°C for imaging. Primary antibody used were: anti-FasII 1/50 (ID4 Hybridoma Bank), anti-GFP 1/500 (Invitrogen A11122), anti-Profilin 1/10 (chi 1J Hybridoma Bank). Secondary antibody used were Alexa Fluor 555 Donkey anti-Mouse, Alexa Fluor 488 Goat Anti-Rabbit at a dilution of 1/500.

### NMJ analysis

For NMJ staining, third instar larvae were dissected in cold PBS and fixed with 4% paraformaldehyde in PBS for 45 min. Larvae were then washed in PBS-T (PBS + 0.5% Triton X-100) six times for 30 min and incubated overnight at 4°C with the following primary antibodies: mouse anti-synaptotagmin, 1:200 (3H2 2D7, Developmental Studies Hybridoma Bank, DSHB) and TRITC-conjugated anti-HRP, 1:1000. After six 30 min washes with PBS-T, secondary antibody anti-mouse conjugated to Alexa-488 was used at a concentration of 1:1000 and incubated at room temperature for 2 h. Larvae were washed again six times with PBS-T and finally mounted in Vectashield.

Images from muscles 6–7 (segment A2-A3) were acquired with a Leica Confocal Microscope SP5. Serial optical sections at 1,024 × 1,024 pixels with 0.4 µM thickness were obtained with the ×40 objective. Bouton number was quantified using Imaris 9 software. ImageJ software was used to measure the muscles area and the NMJ axon length and branching. Statistical tests were performed in GraphPad (PRISM 8).

### RNA probe pulldown

For RNA probe pulldown in BG3 cells, cells were washed with DPBS, harvested and pelleted for 10 minutes at 400g. The cell pellet was re-suspended in lysis buffer (10 mM Tris-HCl at pH 7.4, 150 mM NaCl, 2 mM EDTA, 0.5% NP-40, 0.5 mM DTT, protease inhibitor) and rotated head-over-tail for 15 min at 4°C. Nuclei were collected by 10 min centrifugation at 1000g at 4°C, re-suspended in 300 μl of lysis buffer and sonicated with 5 cycles of 30 s ON, 30 s OFF low power setting. Cytoplasmic and nuclear fractions were joined and centrifuged at 18000g for 10 min at 4°C to remove the remaining cell debris. Protein concentration was determined by Bradford and 1 mg lysate was used for the following pulldown procedure.

For RNA probe pulldown using *E. coli* lysate, *E. coli* cells expressing the indicated proteins were resuspended in lysis buffer (2% TritonX, 20 mM Tris-HCl, 10 mM EDTA, 120 mM NaCl, 50 mM KCl, 1 mM DTT, protease inhibitor) and lysed by sonification. Cell debris and insoluble proteins were removed by centrifugation for 10 min at 12000g at 4°C. The protein concentration was determined by Bradford and the indicated lysate amounts were used for the pulldown procedure.

For RNA probe pulldown using purified recombinant proteins, the indicated purified proteins were resuspended in binding buffer (2% TritonX, 20 mM Tris-HCl, 10 mM EDTA, 120 mM NaCl, 50 mM KCl, 1 mM DTT, protease inhibitor) and used for the following pulldown procedure.

The lysate and purified proteins were incubated with 3 µg of biotinylated RNA probe coupled to 20 µl Dynabeads MyOne Streptavidin C1 (Thermo Fisher) in 600 µl lysis buffer at 4°C for 1 hour rotating head-over-tail. The beads were washed three times with lysis buffer and bound proteins eluted by incubation in 1x Nupage LDS supplemented with 100 mM DTT for 10 minutes at 70°C. Eluted proteins were analyzed by PAGE followed by Coomassie staining, immunoblotting using the corresponding antibodies or proceeded to quantitative proteomic analysis.

### Expression and purification of recombinant proteins

N-His_6_-tagged Fmr1 (220-684) and N-GST-tagged Ythdf (full length) were expressed from pQIq-His6 and pGEX-6P-1, respectively in *E. coli* Rosetta™ 2 (DE3) pLysS (Novagen). Cells were grown in LB-Luria at 37□°C and 160□rpm to an OD_600_ of 0.6–0.8, chilled on ice and expression was induced by addition of IPTG (1□mM). Cells were further incubated at 17□°C and 160□rpm for 20□hours, harvested by centrifugation for 15 min at 4000g at 4°C and snap frozen in liquid nitrogen.

For recombinant protein purification, cells were resuspended in lysis buffer (<u>Hispurification</u>: 25□mM Tris-HCl pH□8, 300□mM NaCl, 20□mM imidazole, 5% glycerol, 1□mM DTT, 1□mM MgCl_2_, Benzonase 1:1000, Protease inhibitors; <u>GST purification:</u> PBS, 5% glycerol, 1mM DTT, 1mM MgCl2, Benzonase 1:1000, Protease inhibitors) and lysed by sonification. Lysates were cleared by centrifugation (45,000×*g*, 30□min, 4□°C). Purification steps were carried out using a Biorad NGC Quest FPLC system. Proteins were passed over an either HisTrap or GstTrap FF 5□ml column (GE Healthcare), respectively. After extensive washing with lysis buffer His-tagged proteins were eluted using lysis buffer containing 500□mM imidazole and GST-tagged proteins were eluted using lysis buffer containing 50 mM reduced Glutathion. Elution fractions were pooled and diluted 1:10 with heparin-binding buffer (0.5X PBS; pH□7.4, 5% glycerol, 1□mM DTT). Diluted proteins were subsequently passed over a HiTrap Heparin HP 5□ml column (GE Healthcare). After washing with binding buffer, recombinant proteins were eluted by running a linear gradient of 0–1.5□M NaCl in binding buffer over 20 column volumes. Fractions containing the respective recombinant proteins were pooled and concentrated using Amicon® Ultra-15 spin concentrators with 10 kDa cut-off (Merck Millipore). Concentrated protein pools were run on a Superdex 200 Increase 10/300 GL, or Superdex 200 16/60 pg in gel filtration buffer (PBS, 1□mM DTT, 10% glycerol). Peak fractions containing the recombinant proteins were pooled, aliquoted, and snap-frozen in liquid nitrogen. Frozen aliquots were stored at −□80□°C.

### TRIBE

TRIBE was performed as previously described (McMahon et al., 2016; Worpenberg et al., 2019; Xu et al., 2018). Briefly, for the identification of mRNA targets in S2R+ cells, Flag-tagged Ythdf and Fmr1 versions fused to the catalytic domain of Adar (cdAdar) were ectopically expressed using a metal inducible expression system. 48 hours after protein expression induction, the cells were washed with DPBS and harvested.

For the identification of in vivo targets, 3^rd^ instar larvae expressing a Flag-tagged version of Ythdf fused to cdAdar driven by *Elav-Gal4* were collected and the brains dissected and collected in PBS.

For protein expression analysis, 50% of the cells/brains were re-suspended in lysis buffer (50 mM Tris-HCl at pH 7.4, 150 mM NaCl, 0.5% NP-40, protease inhibitor) and incubated for 30 minutes on ice. Cell debris was removed by centrifugation for 5 min at 12000 g at 4°C and the expression analyzed by immunoblotting using anti-Flag antibody. For the identification of editing events, the remaining 50% of cells were used for total RNA isolation using Trizol reagent, mRNA was purified by two rounds of Oligo(dT) selection using Dynabeads and the purified mRNA was used for Illumina Next-generation sequencing library preparation using NEBNext^®^ Ultra^™^ II RNA Library Prep Kit for Illumina according to manufacture protocol.

### Co-Immunoprecipitation

S2R+ cells were transfected using Effectene transfection reagent with plasmids expressing the indicated constructs. After 72 hours the cells were washed with DPBS, harvested and pelleted by centrifugation for 3 minutes at 1000 g. The cell pellet was re-suspended in 1ml lysis buffer (50 mM Tris-HCl at pH 7.4, 150 mM NaCl, 0.5% NP-40) supplemented with protease inhibitors and incubated for 30 minutes on ice. The cell debris was removed by centrifugation at 12000 g for 5 minutes at 4°C and the protein concentration of the cleared lysate was measured by Bradford. 2 mg protein lysate was combined with 2 μl antibody coupled to 20 μl Protein G Dynabeads in lysis buffer and incubated rotating head-over-end at 4°C for 2 hours. The beads were washed three times for 10 minutes with lysis buffer and the proteins eluted in 1x NuPage LDS buffer supplemented with 100 mM DTT by incubation for 10 min at 70°C. The eluted proteins were either analyzed by Western Blot or conducted to quantitative proteomics analysis.

### RNA immunoprecipitations (RIP)

For RIP from cell lysate, S2R+ cells were transfected with the plasmids expressing the indicated constructs. After 72 hours the cells were washed with DPBS, harvested and pelleted by centrifugation for 3 minutes at 1000 g. The cell pellet was re-suspended in 1 ml lysis buffer (50 mM Tris-HCl at pH 7.4, 150 mM NaCl, 0.5% NP-40) supplemented with protease inhibitors and RNase inhibitor.

For RIP from *Drosophila* larvae, 30 larvae with the indicated genotype were collected in DPBS on ice, carefully washed by exchanging the DPBS two times and lysed by squishing with a pestle in 1 ml lysis buffer. The resuspended cell pellet or larvae lysate was incubated for 30 minutes on ice. The cell debris was removed by centrifugation at 12000 g for 5 minutes at 4°C and the protein concentration of the cleared lysate was measured by Bradford. 2 mg protein lysate was combined with 2 μl antibody coupled to 20 ul Protein G Dynabeads in lysis buffer and incubated rotating head-over-end at 4°Cfor 2 hours. The beads were washed three times for 10 minutes with lysis buffer. 20% of the beads were used for the elution of protein-RNA complexes by incubation in 1x Nupage LDS buffer supplemented with 100 mM DTT for 10 min at 70°C. The eluted proteins were analyzed by Western Blot. 80% of the beads were used to extract RNA using Trizol reagent. The enrichment of the transcripts was analyzed by qPCR and the fold change calculated by normalizing the transcript levels in the pulldown fractions to the corresponding input and a control pulldown.

For RIP from pupal heads for endogenous Fmr1: 100 heads per each genotype were collected in PBS on ice and subsequently re-suspended in 100 μl of Lysis Buffer (10 mM Tris-HCl pH 7.4, 140 mM NaCl, 0.5%Triton X-100, 1 mM DTT, 2 mM EDTA) supplemented with protease, phosphatase and RNase inhibitors. The heads were lysed with a motor pestle and left on ice for 20’. The cell debris was removed by centrifugation at 12000 g for 10 minutes at 4°C, 1/10 of the volume was taken as an input and kept at −80, while the rest of the supernatant was combined with 15 μl of anti–Fmr1 5B6 and 15 μl of anti–Fmr1 5A11 coupled to 50 ul Protein G Dynabeads in lysis buffer. The lysate was incubated with the beads-coupled antibody rotating head-over-end at 4°C over-night. The day after the beads were washed three times for 10 minutes and subsequently resuspended in 100 μl of lysis buffer that were further treated with 40 µg of Proteinase K for 30’ at 55°C. Upon treatment the supernatant was separated from the beads and RNA was extracted using Trizol reagent. The enrichment of transcripts was analyzed by qPCR and the fold change calculated by normalizing the transcript levels in the pulldown fractions to the corresponding input and a control pulldown. For the RIP with overexpressed Ythdf-HA, 3 µg of anti-HA (Santa Cruz sc-805 rabbit polyclonal) were conjugated to 50 µl of Protein G Dynabeads and used to immunoprecipitate tagged Ythdf from 50 adult heads. For the RIP with overexpressed Fmr1-GFP, 25 µl of GFP Trap_MA beads (Chromtek) were used to immunoprecipitate tagged Fmr1 from 50 adult heads.

### Cell culture

*Drosophila melanogaster* S2R+ and BG3 cells were cultured in Schneider’s *Drosophila* media (PAN BIOTECH) supplemented with 10% FBS and 1% penicillin-streptomycin. The culture medium for BG3 cells was additionally supplemented with 10 µg/ml insulin. Plasmid transfections were achieved using Effectene transfection reagent (Qiagen) according to the manufacture’s protocol.

### RNA isolation & quantitative real-time PCR analysis

Total RNA from S2R+ cells or larval brains was isolated using Trizol reagent according to the manufacturer protocol, DNA contamination removed by DNaseI (NEB) treatment and reverse transcribed using M-MLV reverse transcriptase (Promega). The transcript levels were quantified by real-time PCR using Power SYBR green Master Mix. *Rpl15* mRNA and *18S* rRNA were used as housekeeping control genes.

Total RNA from head lysates was isolated using Trizol reagent according to the manufacturer protocol and reverse transcribed using RevertAid First Strand cDNA Synthesis Kit. The transcript levels were quantified by real-time PCR using Power SYBR green Master Mix. *RP49* and *Tubulin* RNA were used as housekeeping control genes.

### Western Blot analysis

Proteins were extracted for 30 min on ice, the lysates were centrifuged at 12,000*g* for 5 min at 4°C, and protein concentration in the supernatant was determined by Bradford. For Futsch protein detection, lysates were additionally dephosphorylated using Lambda protein phosphatase (NEB) following manufacturer’s instructions. Protein samples were separated on SDS-PAGE gels. Wet transfer to nitrocellulose membrane (Whatman) was performed for 90 min at 100 V. Membranes were blocked for 30 min in 5% nonfat dry milk and PBS–0.5% Tween 20 (PBST) and incubated with primary antibodies overnight at 4°C: anti-Profilin chi1J 1/500 (Hybridoma Bank), anti-Actin I-19 (Santa Cruz sc-1616), anti-Futsch 22C10 (Hybridoma Bank). Signal was detected with corresponding HRP-conjugated secondary antibodies and ECL™ Prime Western Blotting Detection Reagent or ECL™ Select Western Blotting Detection Reagent (Amersham).

### GST-pulldown

Indicated amounts of purified recombinant GST-tagged Ythdf or GST were mixed with purified His6-tagged N-terminally truncated Fmr1 in 1 ml binding buffer (PBS, 5% glycerol, 2% TritonX, 1 mM DTT) and incubated for 1 hour rotating head over end at 4°C with 20 µl Glutathione Sepharose beads. The beads were washed three times with lysis buffer and bound proteins eluted by incubation in 1x Nupage LDS supplemented with 100 mM DTT for 10 minutes at 70°C. The pulled down proteins were analyzed by SDS-PAGE.

### TRAP

TRAP was performed as previously described (Thomas et al., 2012). Briefly, *Drosophila melanogaster* 3^rd^ instar larvae expressing *Elav*-Gal4 driven GFP-tagged Rpl10Ab or cytoplasmic GFP in a wildtype and *Ythdf* mutant background were collected and lysed in extraction buffer (20 mM HEPES, pH 7.5, 150 mM KCl, 5 mM MgCl_2_, 1% Triton X-100, 0.5 mM DTT, 100 µg/mL Cyclohexamide, 100 U/mL Rnase inhibitor, Protease Inhibitor). Cell debris was removed by centrifugation at 12000 g for 5 minutes at 4°C and the protein amount determined by Bradford. 2 mg lysate was combined with 20 μl Protein G Dynabeads conjugated to anti-GFP antibody and incubated at 4°C for two hours followed by three washing steps in Wash Buffer (150 mM NaCl, 0.05% Triton X-100, 50 mM Tris, 5 mM MgCl_2_, and 40 U/mL RNase inhibitor) at 4°C. RNA was extracted using Trizol reagent following manufacture’s protocol and the enrichment of the transcripts analyzed by qPCR.

### m^6^A-miCLIP

miCLIP was performed following previously described method (Linder et al., 2015) using 10 μg of purified mRNA from *Drosophila* S2R+ cells and 5 μg of anti-m^6^A antibody (Synaptic Systems, Lot# 202003/2-82). Immunoprecipitations were performed in quadruplicates and as a control one immunoprecipitation was performed where UV-crosslinking was ommited. Of note, this sample produced a library of limited complexity, reflecting a low amount of background mRNA binding. Briefly, total RNA was isolated using Trizol reagent (Invitrogen) and DNA was removed with DNase-I treatment (NEB). polyadenylated RNA was purified by two rounds of binding to Oligo (dT)25 magnetic beads (NEB) and mRNA was fragmented with RNA fragmentation solution (Ambion) using 1 μL of solution per 2 μg of mRNA and with 7 min incubation at 70°C. Immunoprecipitation was performed at 4°C in 500 μL of binding buffer (BB) (50 mM Tris HCl pH 7,4, 150 mM NaCl, 0,5 % NP-40). First, isolated mRNA and antibody were incubated for 2 hours. Samples were then transferred to individual well of a 12 well cell culture plate and crosslinked on ice (two-times at 150 mJ/cm2). Next, 60 μL of magnetic ProteinG beads (Invitrogen) were resuspended in 500 μL of BB and added to the IP sample. Samples were then incubated for additional 2 hours at 4 °C, before washing with ice-cold solutions was performed: 1x with BB, 2x with high salt buffer (50 mM Tris HCl pH 7,4, 1 M NaCl, 1% NP-40, 0,1% SDS), 1x BB, 2x with PNK buffer (20 mM Tris HCl pH 7,4, 10 mM MgCl2, 0,2% Tween). All washes were performed by gentle pipetting and with 1min incubation on ice. Washes with HSB were additionally rotated for 2 min at 4°C. Finally, beads were resuspended in 900 μL of PNK buffer. 40 μL were used for WB analysis to evaluate immunoprecipitation efficiency. Remaining 860 μL were used for library preparation. All steps of library preparation were performed as previously described in (Sutandy et al., 2016). Libraries were sequenced on an Illumina NextSeq500.

For the miCLIP fragmented input-control library, fragmented mRNA, that was also used for miCLIP IP, was first purified using the 1.8X volume of RNAClean XP beads (Beckman Coulter). Following the 20 min incubation at RT, captured RNA was washed 3x with 80% EtOH and eluted in 20 uL of RNase-free water. The library was prepared using ∼50 ng of cleaned, fragmented mRNA using the NEBNext Ultra Directional RNA Library Prep Kit for Illumina (NEB), by omitting the RNA fragmentation step and following the manufacturer’s protocol. For library amplification, 11 PCR cycles were used and indicated primer and adaptor sequences: NEBNext Index 27 Primer for Illumina: 5′-CAAGCAGAAGACGGCATACGAGATAAAGGAATGTGACTGGAGTTCAGACGTG TGCTCTTCCGATC-s-T-3′ (Expected index read: ATTCCT), NEBNext Adaptor for Illumina: 5′-/5Phos/GAT CGG AAG AGC ACA CGT CTG AAC TCC AGT CUA CAC TCT TTC CCT ACA CGA CGC TCT TCC GAT C-s-T-3′. Libraries were sequenced on an Illumina NextSeq500.

### m^6^A-miCLIP analysis

Sequencing qualities were checked for all reads using FastQC (version 0.11.5) (https://www.bioinformatics.babraham.ac.uk/projects/fastqc/). Afterwards, reads were filtered based on sequencing qualities (Phred score) of the barcode region. Reads with more than one position with a Phred score < 20 in the experimental barcode (positions 4 to 7 in the reads) or any position with a Phred score < 10 in the random barcode (positions 1 to 3 and 8 to 9) were excluded from the subsequent analysis. Remaining reads were de-multiplexed based on the experimental barcode (positions 4 to 7) using Flexbar (v3.0.0) (Dodt et al., 2012) without allowing any mismatch.

Individual samples were processed using the CLIP Tool Kit (CTK) v1.0.9. (Shah et al., 2017). We largely followed recommended user guide lines specific to CTK iCLIP data analysis as described here (https://zhanglab.c2b2.columbia.edu/index.php/ICLIP_data_analysis_using_CTK). Briefly, 3’adapter sequences [AGATCGGAAGAGCGGTTCAG] were trimmed using cutadapt v1.8. [overlap□=□5; -m 29] (Martin, 2011). PCR duplicates were removed using a custom perl script, followed by the extraction of the 9 nucleotide miCLIP barcode and its addition to the read name. All cDNA libraries were filtered for common *Drosophila* virus sequences (Webster et al., 2016) using bowtie v1.1.2 [-p 4 -q (-X 1000) --fr best]. Next, to avoid sequencing read alignment software biases, we decided to map sequencing reads to the *Drosophila melanogaster* dm6 genome assembly (ensemble v81) using novoalign (http://www.novocraft.com/), bwa (Li and Durbin, 2009) and STAR v2.4.2a (Dobin et al., 2013). For STAR alignments, we used a custom python script to transform sam files into the expected format for downstream CITS identification. For STAR alignments, we did not consider spliced reads, soft-clipped reads, mismatches and indels near read start and read end, and reads with more than one indel or mismatch. Then, unique tags were identified using *parseAlignment*.*pl* [-v --map-qual 1 --min-len 18 --indel-to-end 2] to extract unique tags, followed by read collapsing using *tag2collapse.pl*. [-v -big --random-barcode -EM 30 --seq-error-model alignment -weight --weight-in-name --keep-max-score --keep-tag-name]. Crosslinking induced mutation sites (CIMS) indicative for the antibody-m^6^A interaction were identified running *CIMS.pl* [-big -n 10], and CIMS with FDR < 0.001 were retained. Crosslinking induced truncations sites were identified using *CITS.pl* [-big -p 0.001 --gap 25]. Sites spanning more than 1 nucleotide were removed.

We further filtered identified CIMS and CITS to be reproducible in at least 2 out of 4 replicate m^6^A-immunoprecipitation samples and not present in identified CITS from the input control sample for each aligner separately. Moreover, CIMS were filtered to have a minimum of 6 unique tags [k > 5], at least three unique substitutions [m > 2], and be prevalent in less than 20% of the coverage [m/k < 0.2] to avoid calling homozygous and heterozygous single nucleotide variants. CIMS sites were found to be almost exclusive C-to-T conversions (n = 6225, 88% ± 5.8%) independent of the alignment software used (3 aligners n = 2677, 2 n = 2411, 1 n = 1137). For a stringent CITS set, we filtered CITS sites (n = 22917) that mostly truncated at A residues (n = 11897, 52%), and were followed by C residues (CITS-AC; n = 6799, 57%). Two thousand three hundred and two (37%) of C-to-T CIMS overlapped within a 1nt window to CITS-AC sites, suggesting that in many cases the same nucleotide was identified. Together, we considered a set of 13024 C-to-T conversions CIMS and AC truncation CITS across 2464 genes for our final S2 cell miCLIP data set. CITS site were annotated as described before (Wessels et al., 2019). For representation purposes we simplified the annotation categories. All CITS not annotated to 5’UTR, CDS, 3’UTR or intron were summarized in the category ‘other’. Enrichments were calculated relative to median feature proportions (5’UTR = 0.08 (131nt), CDS = 0.78 (1309.5nt), 3’UTR = 0.14 (234nt)) determined previously for S2 cells (Wessels et al., 2019). Enrichments for sites annotated as intronic was set to 1.

### SELECT

SELECT was performed as described previously (Mao et al., 2019). Briefly, 1 μg total RNA of isolated 3^rd^ instar larval brains of the indicated genotypes was diluted in 5 μM dNTPs, 40 nM up and 40 nM down primers (m6A_1 up: TAGCCAGTACCGTAGTGCGTGTTGGTTTTGGTTTGGGTG; m6A_1 down: CTTTCTTTGGTTTTGGTTAATAACTCAGAGGCTGAGTCGCTGCAT; m6A_2 up: TAGCCAGTACCGTAGTGCGTGGTTTTTTTTCGACTTTGCTTAATTG; m6A_2 down: CTTTCGGTTTTTGGTTTTGTTCAGAGGCTGAGTCGCTGCAT; Control up: TAGCCAGTACCGTAGTGCGTGGTTTTTTTTCGACTTTGCTTAATTGT; Control down: TTTCGGTTTTTGGTTTTGTTTCAGAGGCTGAGTCGCTGCAT), targeting the m^6^A or a control site one nucleotide upstream of the m^6^A_2 site on *futsch*, and 1× CutSmart buffer (NEB) to 17 μl. Annealing of primers was performed for 1 minute at each 90°C, 80°C, 70°C, 60°C, 50°C and 6 min at 40°C. Afterwards, 0.01U Bst 2.0 DNA polymerase, 0.5U SplintR ligase and 10 nmol ATP was added in a total volume of 3 μl and incubated for 20 minutes at 40°C and 80°C. qPCR for quantification was carried out using 4 ul of SELECT reaction in a 20 μl reaction volume using SYBR green. Relative SELECT products were calculated by normalization to the RNA abundance determined by the control site and the wildtype control.

## Acknowledgments

We thank the Bloomington *Drosophila* Stock Center, the Vienna *Drosophila* Resource Center and the *Drosophila* Genomics Resource Center at Indiana University for stocks, plasmids and cell lines; members of the Quattrone and Roignant labs for helpful discussion. We thank the IMB Genomics core facility for their helpful support and the use of its NextSeq500 (INST 247/870-1 FUGG), and the advanced Imaging Facility at Department CIBIO for great support. Support by IMB Proteomics Core Facility is gratefully acknowledged (instrument is funded by DFG INST 247/766-1 FUGG). In particular, we wish to thank Anja Freiwald from IMB Proteomics core facility for sample preparation and Dr. Mario Dejung from Proteomics core facility for data processing. We thank Prof. Bassem Hassan for kindly sharing *Drosophila* stocks and Dr. Fabian Feiguin for sharing stocks and anti-Futsch antibody. We thank Tobias Jakobi for help with loading TRIBE datasets. Research in the laboratory of J.-Y.R. is supported by the Deutsch-Israelische Projektkooperation (DIP) RO 4681/6-1, the Deutsche Forschungsgemeinschaft RO 4681/9-1 and the Epitran COST action (CA16120). Research in the laboratory of A. Q. is supported by the AIRC Foundation, The CARITRO Foundation, a private donation by the Zobele family and the Epitran COST action (CA16120). The Vermeulen lab is part of the Oncode Institute, which is partly funded by the Dutch Cancer Society (KWF). Alessia Soldano in the A.Q. lab is supported by funding from the European Union’s Horizon 2020 research and Innovation programme under the Marie Skłodowska Curie grant agreement No 752621. Chiara Paolantoni in the lab of J.Y.R. is supported by a Boehringer Ingelheim Fonds fellowship.

## Author contributions

A.S., L.W., A.Q. and J.-Y.R. conceived the study. A.S, L.W., C.P., S.L., M.M., T.L., H.-H.W., M.N., G.A., M.S., R.R.E., A.B., M.M.M., M.V., F.B., U.O., C.D., A.Q., and J.-Y.R. performed the methodology. A.S. and L.W. wrote the draft of the manuscript. All authors reviewed and edited the manuscript. A.S., A.Q. and J.-Y.R. supervised the study.

## References

Angelova, M.T., Dimitrova, D.G., Dinges, N., Lence, T., Worpenberg, L., Carre, C., and Roignant, J.Y. (2018). The Emerging Field of Epitranscriptomics in Neurodevelopmental and Neuronal Disorders. Front Bioeng Biotechnol 6, 46.

Arguello, A.E., DeLiberto, A.N., and Kleiner, R.E. (2017). RNA Chemical Proteomics Reveals the N(6)-Methyladenosine (m(6)A)-Regulated Protein-RNA Interactome. J Am Chem Soc 139, 17249–17252.

Bagni, C., and Zukin, R.S. (2019). A Synaptic Perspective of Fragile X Syndrome and Autism Spectrum Disorders. Neuron 101, 1070–1088.

Bolduc, F.V., Bell, K., Cox, H., Broadie, K.S., and Tully, T. (2008). Excess protein synthesis in Drosophila fragile X mutants impairs long-term memory. Nat Neurosci 11, 1143–1145.

Chang, M., Lv, H., Zhang, W., Ma, C., He, X., Zhao, S., Zhang, Z.W., Zeng, Y.X., Song, S., Niu, Y., et al. (2017). Region-specific RNA m(6)A methylation represents a new layer of control in the gene regulatory network in the mouse brain. Open Biol 7.

Chen, E., Sharma, M.R., Shi, X., Agrawal, R.K., and Joseph, S. (2014). Fragile X mental retardation protein regulates translation by binding directly to the ribosome. Mol Cell 54, 407–417.

Chen, J., Zhang, Y.C., Huang, C., Shen, H., Sun, B., Cheng, X., Zhang, Y.J., Yang, Y.G., Shu, Q., Yang, Y., et al. (2019). m(6)A Regulates Neurogenesis and Neuronal Development by Modulating Histone Methyltransferase Ezh2. Genomics Proteomics Bioinformatics 17, 154–168.

Coffee, R.L., Jr., Tessier, C.R., Woodruff, E.A., 3rd, and Broadie, K. (2010). Fragile X mental retardation protein has a unique, evolutionarily conserved neuronal function not shared with FXR1P or FXR2P. Dis Model Mech 3, 471–485.

Dahlhaus, R. (2018). Of Men and Mice: Modeling the Fragile X Syndrome. Front Mol Neurosci 11, 41.

Darnell, J.C., Van Driesche, S.J., Zhang, C., Hung, K.Y., Mele, A., Fraser, C.E., Stone, E.F., Chen, C., Fak, J.J., Chi, S.W., et al. (2011). FMRP stalls ribosomal translocation on mRNAs linked to synaptic function and autism. Cell 146, 247–261.

Davis, J.K., and Broadie, K. (2017). Multifarious Functions of the Fragile X Mental Retardation Protein. Trends Genet 33, 703–714.

Dobin, A., Davis, C.A., Schlesinger, F., Drenkow, J., Zaleski, C., Jha, S., Batut, P., Chaisson, M., and Gingeras, T.R. (2013). STAR: ultrafast universal RNA-seq aligner. Bioinformatics 29, 15–21.

Dodt, M., Roehr, J.T., Ahmed, R., and Dieterich, C. (2012). FLEXBAR-Flexible Barcode and Adapter Processing for Next-Generation Sequencing Platforms. Biology (Basel) 1, 895–905.

Dominissini, D., Moshitch-Moshkovitz, S., Schwartz, S., Salmon-Divon, M., Ungar, L., Osenberg, S., Cesarkas, K., Jacob-Hirsch, J., Amariglio, N., Kupiec, M., et al. (2012). Topology of the human and mouse m6A RNA methylomes revealed by m6A-seq. Nature 485, 201–206.

Drozd, M., Bardoni, B., and Capovilla, M. (2018). Modeling Fragile X Syndrome in Drosophila. Front Mol Neurosci 11, 124.

Du, K., Zhang, L., Lee, T., and Sun, T. (2019). m(6)A RNA Methylation Controls Neural Development and Is Involved in Human Diseases. Mol Neurobiol 56, 1596–1606.

Edens, B.M., Vissers, C., Su, J., Arumugam, S., Xu, Z., Shi, H., Miller, N., Rojas Ringeling, F., Ming, G.L., He, C., et al. (2019). FMRP Modulates Neural Differentiation through m(6)A-Dependent mRNA Nuclear Export. Cell Rep 28, 845–854 e845.

Edupuganti, R.R., Geiger, S., Lindeboom, R.G.H., Shi, H., Hsu, P.J., Lu, Z., Wang, S.Y., Baltissen, M.P.A., Jansen, P., Rossa, M., et al. (2017). N(6)-methyladenosine (m(6)A) recruits and repels proteins to regulate mRNA homeostasis. Nat Struct Mol Biol 24, 870–878.

Engel, M., Eggert, C., Kaplick, P.M., Eder, M., Roh, S., Tietze, L., Namendorf, C., Arloth, J., Weber, P., Rex-Haffner, M., et al. (2018). The Role of m(6)A/m-RNA Methylation in Stress Response Regulation. Neuron 99, 389–403 e389.

Garber, K., Smith, K.T., Reines, D., and Warren, S.T. (2006). Transcription, translation and fragile X syndrome. Curr Opin Genet Dev 16, 270–275.

Garber, K.B., Visootsak, J., and Warren, S.T. (2008). Fragile X syndrome. Eur J Hum Genet 16, 666–672.

Garcia-Campos, M.A., Edelheit, S., Toth, U., Safra, M., Shachar, R., Viukov, S., Winkler, R., Nir, R., Lasman, L., Brandis, A., et al. (2019). Deciphering the “m(6)A Code” via Antibody-Independent Quantitative Profiling. Cell 178, 731–747 e716.

Hagerman, R.J., Des-Portes, V., Gasparini, F., Jacquemont, S., and Gomez-Mancilla, B. (2014). Translating molecular advances in fragile X syndrome into therapy: a review. J Clin Psychiatry 75, e294–307.

Hagerman, R.J., Miller, L.J., McGrath-Clarke, J., Riley, K., Goldson, E., Harris, S.W., Simon, J., Church, K., Bonnell, J., Ognibene, T.C., et al. (2002). Influence of stimulants on electrodermal studies in Fragile X syndrome. Microsc Res Tech 57, 168–173.

Haussmann, I.U., Bodi, Z., Sanchez-Moran, E., Mongan, N.P., Archer, N., Fray, R.G., and Soller, M. (2016). m(6)A potentiates Sxl alternative pre-mRNA splicing for robust Drosophila sex determination. Nature 540, 301–304.

Hsu, P.J., Shi, H., Zhu, A.C., Lu, Z., Miller, N., Edens, B.M., Ma, Y.C., and He, C. (2019). The RNA-binding protein FMRP facilitates the nuclear export of N (6)-methyladenosine-containing mRNAs. J Biol Chem 294, 19889–19895.

Ishizuka, A., Siomi, M.C., and Siomi, H. (2002). A Drosophila fragile X protein interacts with components of RNAi and ribosomal proteins. Genes Dev 16, 2497–2508.

Jacquemont, S., Pacini, L., Jonch, A.E., Cencelli, G., Rozenberg, I., He, Y., D’Andrea, L., Pedini, G., Eldeeb, M., Willemsen, R., et al. (2018). Protein synthesis levels are increased in a subset of individuals with fragile X syndrome. Hum Mol Genet 27, 2039–2051.

Jaenisch, R., and Bird, A. (2003). Epigenetic regulation of gene expression: how the genome integrates intrinsic and environmental signals. Nat Genet 33 *Suppl*, 245–254.

Jia, G., Fu, Y., Zhao, X., Dai, Q., Zheng, G., Yang, Y., Yi, C., Lindahl, T., Pan, T., Yang, Y.G., et al. (2011). N6-methyladenosine in nuclear RNA is a major substrate of the obesity-associated FTO. Nat Chem Biol 7, 885–887.

Jung, Y., and Goldman, D. (2018). Role of RNA modifications in brain and behavior. Genes Brain Behav 17, e12444.

Kan, L., Grozhik, A.V., Vedanayagam, J., Patil, D.P., Pang, N., Lim, K.S., Huang, Y.C., Joseph, B., Lin, C.J., Despic, V., et al. (2017). The m(6)A pathway facilitates sex determination in Drosophila. Nat Commun 8, 15737.

Kashima, R., Redmond, P.L., Ghatpande, P., Roy, S., Kornberg, T.B., Hanke, T., Knapp, S., Lagna, G., and Hata, A. (2017). Hyperactive locomotion in a Drosophila model is a functional readout for the synaptic abnormalities underlying fragile X syndrome. Sci Signal 10.

Kidd, S.A., Lachiewicz, A., Barbouth, D., Blitz, R.K., Delahunty, C., McBrien, D., Visootsak, J., and Berry-Kravis, E. (2014). Fragile X syndrome: a review of associated medical problems. Pediatrics 134, 995–1005.

Klemmer, P., Meredith, R.M., Holmgren, C.D., Klychnikov, O.I., Stahl-Zeng, J., Loos, M., van der Schors, R.C., Wortel, J., de Wit, H., Spijker, S., et al. (2011). Proteomics, ultrastructure, and physiology of hippocampal synapses in a fragile X syndrome mouse model reveal presynaptic phenotype. J Biol Chem 286, 25495–25504.

Koranda, J.L., Dore, L., Shi, H., Patel, M.J., Vaasjo, L.O., Rao, M.N., Chen, K., Lu, Z., Yi, Y., Chi, W., et al. (2018). Mettl14 Is Essential for Epitranscriptomic Regulation of Striatal Function and Learning. Neuron 99, 283–292 e285.

Laggerbauer, B., Ostareck, D., Keidel, E.M., Ostareck-Lederer, A., and Fischer, U. (2001). Evidence that fragile X mental retardation protein is a negative regulator of translation. Hum Mol Genet 10, 329–338.

Lence, T., Akhtar, J., Bayer, M., Schmid, K., Spindler, L., Ho, C.H., Kreim, N., Andrade-Navarro, M.A., Poeck, B., Helm, M., et al. (2016). m(6)A modulates neuronal functions and sex determination in Drosophila. Nature 540, 242–247.

Lence, T., Paolantoni, C., Worpenberg, L., and Roignant, J.Y. (2019). Mechanistic insights into m(6)A RNA enzymes. Biochim Biophys Acta Gene Regul Mech 1862, 222–229.

Lence, T., Soller, M., and Roignant, J.Y. (2017). A fly view on the roles and mechanisms of the m(6)A mRNA modification and its players. RNA Biol 14, 1232–1240.

Li, H., and Durbin, R. (2009). Fast and accurate short read alignment with Burrows-Wheeler transform. Bioinformatics 25, 1754–1760.

Li, J., Yang, X., Qi, Z., Sang, Y., Liu, Y., Xu, B., Liu, W., Xu, Z., and Deng, Y. (2019). The role of mRNA m(6)A methylation in the nervous system. Cell Biosci 9, 66.

Li, L., Zang, L., Zhang, F., Chen, J., Shen, H., Shu, L., Liang, F., Feng, C., Chen, D., Tao, H., et al. (2017). Fat mass and obesity-associated (FTO) protein regulates adult neurogenesis. Hum Mol Genet 26, 2398–2411.

Li, M., Zhao, X., Wang, W., Shi, H., Pan, Q., Lu, Z., Perez, S.P., Suganthan, R., He, C., Bjoras, M., et al. (2018). Ythdf2-mediated m(6)A mRNA clearance modulates neural development in mice. Genome Biol 19, 69.

Li, Z., Zhang, Y., Ku, L., Wilkinson, K.D., Warren, S.T., and Feng, Y. (2001). The fragile X mental retardation protein inhibits translation via interacting with mRNA. Nucleic Acids Res 29, 2276–2283.

Liao, S., Sun, H., and Xu, C. (2018). YTH Domain: A Family of N(6)-methyladenosine (m(6)A) Readers. Genomics Proteomics Bioinformatics 16, 99–107.

Linder, B., Grozhik, A.V., Olarerin-George, A.O., Meydan, C., Mason, C.E., and Jaffrey, S.R. (2015). Single-nucleotide-resolution mapping of m6A and m6Am throughout the transcriptome. Nat Methods 12, 767–772.

Livneh, I., Moshitch-Moshkovitz, S., Amariglio, N., Rechavi, G., and Dominissini, D. (2020). The m(6)A epitranscriptome: transcriptome plasticity in brain development and function. Nat Rev Neurosci 21, 36–51.

Lu, R., Wang, H., Liang, Z., Ku, L., O’Donnell W, T., Li, W., Warren, S.T., and Feng, Y. (2004). The fragile X protein controls microtubule-associated protein 1B translation and microtubule stability in brain neuron development. Proc Natl Acad Sci U S A 101, 15201–15206.

Ma, C., Chang, M., Lv, H., Zhang, Z.W., Zhang, W., He, X., Wu, G., Zhao, S., Zhang, Y., Wang, D., et al. (2018). RNA m(6)A methylation participates in regulation of postnatal development of the mouse cerebellum. Genome Biol 19, 68.

Mao, Y., Dong, L., Liu, X.M., Guo, J., Ma, H., Shen, B., and Qian, S.B. (2019). m(6)A in mRNA coding regions promotes translation via the RNA helicase-containing YTHDC2. Nat Commun 10, 5332.

Martin, M. (2011). Cutadapt Removes Adapter Sequences from High-Throughput Sequencing Reads. EMBnet Journal 17, 10–12.

Maurin, T., Zongaro, S., and Bardoni, B. (2014). Fragile X Syndrome: from molecular pathology to therapy. Neurosci Biobehav Rev 46 *Pt* *2*, 242–255.

McMahon, A.C., Rahman, R., Jin, H., Shen, J.L., Fieldsend, A., Luo, W., and Rosbash, M. (2016). TRIBE: Hijacking an RNA-Editing Enzyme to Identify Cell-Specific Targets of RNA-Binding Proteins. Cell 165, 742–753.

Merkurjev, D., Hong, W.T., Iida, K., Oomoto, I., Goldie, B.J., Yamaguti, H., Ohara, T., Kawaguchi, S.Y., Hirano, T., Martin, K.C., et al. (2018). Synaptic N(6)-methyladenosine (m(6)A) epitranscriptome reveals functional partitioning of localized transcripts. Nat Neurosci 21, 1004–1014.

Meyer, K.D., Saletore, Y., Zumbo, P., Elemento, O., Mason, C.E., and Jaffrey, S.R. (2012). Comprehensive analysis of mRNA methylation reveals enrichment in 3’ UTRs and near stop codons. Cell 149, 1635–1646.

Michel, C.I., Kraft, R., and Restifo, L.L. (2004). Defective neuronal development in the mushroom bodies of Drosophila fragile X mental retardation 1 mutants. J Neurosci 24, 5798–5809.

Patil, D.P., Pickering, B.F., and Jaffrey, S.R. (2018). Reading m(6)A in the Transcriptome: m(6)A-Binding Proteins. Trends Cell Biol 28, 113–127.

Reeve, S.P., Bassetto, L., Genova, G.K., Kleyner, Y., Leyssen, M., Jackson, F.R., and Hassan, B.A. (2005). The Drosophila fragile X mental retardation protein controls actin dynamics by directly regulating profilin in the brain. Curr Biol 15, 1156–1163.

Roignant, J.Y., and Soller, M. (2017). m(6)A in mRNA: An Ancient Mechanism for Fine-Tuning Gene Expression. Trends Genet 33, 380–390.

Rousseau, F., Labelle, Y., Bussieres, J., and Lindsay, C. (2011). The fragile x mental retardation syndrome 20 years after the FMR1 gene discovery: an expanding universe of knowledge. Clin Biochem Rev 32, 135–162.

Sahoo, P.K., Lee, S.J., Jaiswal, P.B., Alber, S., Kar, A.N., Miller-Randolph, S., Taylor, E.E., Smith, T., Singh, B., Ho, T.S., et al. (2018). Axonal G3BP1 stress granule protein limits axonal mRNA translation and nerve regeneration. Nat Commun 9, 3358.

Santoro, M.R., Bray, S.M., and Warren, S.T. (2012). Molecular mechanisms of fragile X syndrome: a twenty-year perspective. Annu Rev Pathol 7, 219–245.

Schaefer, T.L., Davenport, M.H., and Erickson, C.A. (2015). Emerging pharmacologic treatment options for fragile X syndrome. Appl Clin Genet 8, 75–93.

Shah, A., Qian, Y., Weyn-Vanhentenryck, S.M., and Zhang, C. (2017). CLIP Tool Kit (CTK): a flexible and robust pipeline to analyze CLIP sequencing data. Bioinformatics 33, 566–567.

Shi, H., Zhang, X., Weng, Y.L., Lu, Z., Liu, Y., Lu, Z., Li, J., Hao, P., Zhang, Y., Zhang, F., et al. (2018). m(6)A facilitates hippocampus-dependent learning and memory through YTHDF1. Nature 563, 249–253.

Stefani, G., Fraser, C.E., Darnell, J.C., and Darnell, R.B. (2004). Fragile X mental retardation protein is associated with translating polyribosomes in neuronal cells. J Neurosci 24, 7272–7276.

Suhl, J.A., Chopra, P., Anderson, B.R., Bassell, G.J., and Warren, S.T. (2014). Analysis of FMRP mRNA target datasets reveals highly associated mRNAs mediated by G-quadruplex structures formed via clustered WGGA sequences. Hum Mol Genet 23, 5479–5491.

Sutandy, F.X., Hildebrandt, A., and Konig, J. (2016). Profiling the Binding Sites of RNA-Binding Proteins with Nucleotide Resolution Using iCLIP. Methods Mol Biol 1358, 175–195.

Thomas, A., Lee, P.J., Dalton, J.E., Nomie, K.J., Stoica, L., Costa-Mattioli, M., Chang, P., Nuzhdin, S., Arbeitman, M.N., and Dierick, H.A. (2012). A versatile method for cell-specific profiling of translated mRNAs in Drosophila. PLoS One 7, e40276.

Utari, A., Adams, E., Berry-Kravis, E., Chavez, A., Scaggs, F., Ngotran, L., Boyd, A., Hessl, D., Gane, L.W., Tassone, F., et al. (2010). Aging in fragile X syndrome. J Neurodev Disord 2, 70–76.

Verheyen, E.M., and Cooley, L. (1994). Profilin mutations disrupt multiple actin-dependent processes during Drosophila development. Development 120, 717–728.

Wang, C.X., Cui, G.S., Liu, X., Xu, K., Wang, M., Zhang, X.X., Jiang, L.Y., Li, A., Yang, Y., Lai, W.Y., et al. (2018). METTL3-mediated m6A modification is required for cerebellar development. PLoS Biol 16, e2004880.

Wang, X., Zhao, B.S., Roundtree, I.A., Lu, Z., Han, D., Ma, H., Weng, X., Chen, K., Shi, H., and He, C. (2015). N(6)-methyladenosine Modulates Messenger RNA Translation Efficiency. Cell 161, 1388–1399.

Webster, C.L., Longdon, B., Lewis, S.H., and Obbard, D.J. (2016). Twenty-Five New Viruses Associated with the Drosophilidae (Diptera). Evol Bioinform Online 12, 13–25.

Weng, Y.L., Wang, X., An, R., Cassin, J., Vissers, C., Liu, Y., Liu, Y., Xu, T., Wang, X., Wong, S.Z.H., et al. (2018). Epitranscriptomic m(6)A Regulation of Axon Regeneration in the Adult Mammalian Nervous System. Neuron 97, 313–325 e316.

Wessels, H.H., Lebedeva, S., Hirsekorn, A., Wurmus, R., Akalin, A., Mukherjee, N., and Ohler, U. (2019). Global identification of functional microRNA-mRNA interactions in Drosophila. Nat Commun 10, 1626.

Widagdo, J., and Anggono, V. (2018). The m6A-epitranscriptomic signature in neurobiology: from neurodevelopment to brain plasticity. J Neurochem 147, 137–152.

Widagdo, J., Zhao, Q.Y., Kempen, M.J., Tan, M.C., Ratnu, V.S., Wei, W., Leighton, L., Spadaro, P.A., Edson, J., Anggono, V., et al. (2016). Experience-Dependent Accumulation of N6-Methyladenosine in the Prefrontal Cortex Is Associated with Memory Processes in Mice. J Neurosci 36, 6771–6777.

Worpenberg, L., Jakobi, T., Dieterich, C., and Roignant, J.Y. (2019). Identification of Methylated Transcripts Using the TRIBE Approach. Methods Mol Biol 1870, 89–106.

Xiao, Y., Wang, Y., Tang, Q., Wei, L., Zhang, X., and Jia, G. (2018). An Elongation- and Ligation-Based qPCR Amplification Method for the Radiolabeling-Free Detection of Locus-Specific N(6) -Methyladenosine Modification. Angew Chem Int Ed Engl 57, 15995–16000.

Xu, D., Shen, W., Guo, R., Xue, Y., Peng, W., Sima, J., Yang, J., Sharov, A., Srikantan, S., Yang, J., et al. (2013). Top3beta is an RNA topoisomerase that works with fragile X syndrome protein to promote synapse formation. Nat Neurosci 16, 1238–1247.

Xu, W., Rahman, R., and Rosbash, M. (2018). Mechanistic implications of enhanced editing by a HyperTRIBE RNA-binding protein. RNA 24, 173–182.

Yoon, K.J., Ming, G.L., and Song, H. (2018). Epitranscriptomes in the Adult Mammalian Brain: Dynamic Changes Regulate Behavior. Neuron 99, 243–245.

Yu, J., Chen, M., Huang, H., Zhu, J., Song, H., Zhu, J., Park, J., and Ji, S.J. (2018). Dynamic m6A modification regulates local translation of mRNA in axons. Nucleic Acids Res 46, 1412–1423.

Zhang, F., Kang, Y., Wang, M., Li, Y., Xu, T., Yang, W., Song, H., Wu, H., Shu, Q., and Jin, P. (2018). Fragile X mental retardation protein modulates the stability of its m6A-marked messenger RNA targets. Hum Mol Genet 27, 3936–3950.

Zhang, Y.Q., Bailey, A.M., Matthies, H.J., Renden, R.B., Smith, M.A., Speese, S.D., Rubin, G.M., and Broadie, K. (2001). Drosophila fragile X-related gene regulates the MAP1B homolog Futsch to control synaptic structure and function. Cell 107, 591–603.

Zhao, B.S., Roundtree, I.A., and He, C. (2017). Post-transcriptional gene regulation by mRNA modifications. Nat Rev Mol Cell Biol 18, 31–42.

Zhao, Y.L., Liu, Y.H., Wu, R.F., Bi, Z., Yao, Y.X., Liu, Q., Wang, Y.Z., and Wang, X.X. (2019). Understanding m(6)A Function Through Uncovering the Diversity Roles of YTH Domain-Containing Proteins. Mol Biotechnol 61, 355–364.

Zheng, G., Dahl, J.A., Niu, Y., Fedorcsak, P., Huang, C.M., Li, C.J., Vagbo, C.B., Shi, Y., Wang, W.L., Song, S.H., et al. (2013). ALKBH5 is a mammalian RNA demethylase that impacts RNA metabolism and mouse fertility. Mol Cell 49, 18–29.

Zhuang, M., Li, X., Zhu, J., Zhang, J., Niu, F., Liang, F., Chen, M., Li, D., Han, P., and Ji, S.J. (2019). The m6A reader YTHDF1 regulates axon guidance through translational control of Robo3.1 expression. Nucleic Acids Res 47, 4765–4777.

